# Multi-tissue transcriptomic aging atlas reveals predictive aging biomarkers in the killifish

**DOI:** 10.1101/2025.01.28.635350

**Authors:** Emma K. Costa, Jingxun Chen, Ian H. Guldner, Lajoyce Mboning, Natalie Schmahl, Aleksandra Tsenter, Man-Ru Wu, Patricia Moran-Losada, Louis S. Bouchard, Sui Wang, Param Priya Singh, Matteo Pellegrini, Anne Brunet, Tony Wyss-Coray

**Author notes:** These authors contributed equally (Emma K. Costa, Jingxun Chen). These authors contributed equally (Anne Brunet, Tony Wyss-Coray).

## Abstract

Aging is associated with progressive tissue dysfunction, leading to frailty and mortality. Characterizing aging features, such as changes in gene expression and dynamics, shared across tissues or specific to each tissue, is crucial for understanding systemic and local factors contributing to the aging process. We performed RNA-sequencing on 13 tissues at 6 different ages in the African turquoise killifish, the shortest-lived vertebrate that can be raised in captivity. This comprehensive, sex-balanced ‘atlas’ dataset reveals the varying strength of sex-age interactions across killifish tissues and identifies age-altered biological pathways that are evolutionarily conserved. Demonstrating the utility of this resource, we discovered that the killifish head kidney exhibits a myeloid bias during aging, a phenomenon more pronounced in females than in males. In addition, we developed tissue-specific ‘transcriptomic clocks’ and identified biomarkers predictive of chronological age. We show the importance of sex-specific clocks for selected tissues and use the tissue clocks to evaluate a dietary intervention in the killifish. Our work provides a comprehensive resource for studying aging dynamics across tissues in the killifish, a powerful vertebrate aging model.

## Introduction

Aging is the greatest risk factor for disease and death in humans. It is a highly complex process, characterized by progressive cellular and tissue dysfunction. Such dysfunction is accompanied by shared molecular features, referred to as ‘hallmarks of aging’^1^, such as chronic inflammation, loss of proteostasis, and dysregulated nutrient sensing. Recent work in mice suggests that these aging hallmarks can differ between males and females in specific tissues^2–7^. Moreover, the amplitude and the onset age of these hallmarks can also differ among the tissues of an organism^8^. Currently, the extent of sex dimorphism in tissue aging, including age-altered gene pathways and aging trajectories, is not well understood. Understanding the age-sex relationship among diverse tissues will augment our understanding of sex-specific interventions to slow and even reverse aging.

We used the African turquoise killifish (*Nothobranchius furzeri*) as a naturally accelerated vertebrate aging model for our studies. The killifish has emerged as a new vertebrate model in aging research because it has conserved aging signatures and a short lifespan, which are attractive features for rapid lifespan and healthspan intervention testing^9–18^. The median lifespan of the killifish is 4-6 months (about a fifth of the mouse lifespan and a seventh of the zebrafish lifespan), with vertebrate-specific genes, tissues, and systems conserved with humans^9–18^. Several conserved aging mechanisms and interventions have been reported in this model, such as mutants of the nutrient-sensing pathway^19,20^ and the germline^21^, dietary modifications^19,20,22,23^, and administration of small molecule treatments^24–27^. Many of these interventions have sex-specific effects on killifish lifespan, suggesting interesting age-sex relationships in killifish that can provide critical insights into our central question.

Transcriptomic analysis (e.g., RNA-sequencing or single-cell RNA-sequencing) has been applied in the killifish to understand the aging signatures of tissues or cell types and the effects of aging interventions^19–22,26,28–37^. These studies have identified the crucial gene pathways and biological processes altered by tissue aging, such as elevated inflammation^19–21,28,32,34,38^ and loss of proteostasis^28,31,39^. However, publications in killifish have mostly focused on a single tissue or sex and sample only a few time points (2-3 time points), which limits the ability to study gene dynamics across time and tissues. Because direct comparison across multiple tissues is lacking, it remains unknown how similarly the tissue transcriptomes change with age, how biological sex affects the aging pathways in each tissue and across tissues, and which tissues or pathways have early onset of gene expression changes or distinct dynamics with age. A broad characterization of killifish tissue aging will be a valuable resource to pinpoint the specific aspects of vertebrate aging that can be modeled in killifish and are suitable to intervention testing. Such characterization should also allow development of machine-learning models (‘aging clocks’) for rapid evaluation of intervention efficacy.

In this study, we comprehensively profiled the aging transcriptomes of 13 tissues across 6 time points for male and female killifish. This 677-sample dataset is the most comprehensive, high-quality tissue aging atlas of the killifish to date. We identified distinct age-sex relationships for each tissue, the age-correlated genes and pathways shared across multiple tissues, and the tissue-specific genes that may drive cell-type composition changes in the aging head kidney, a main hematopoietic compartment of the killifish. Lastly, we developed tissue-specific aging clocks that allow us to evaluate a published lifespan intervention and to uncover the importance of incorporating sex-specific features in building age prediction models.

## Results

### A large-scale atlas reveals shared and tissue-specific age effects on different tissues

To understand how different tissues age in the killifish, we constructed a multi-tissue transcriptomic aging atlas consisting of 677 samples collected from two independent aging cohorts of killifish (Fig. 1a). We developed a protocol for cardiac perfusion and performed this procedure on these killifish to limit the impact of circulating immune cells on the tissue transcriptome signature, thus allowing discovery of age-dependent changes in tissue-resident cell types. Thirteen tissues (bone, brain, retina/retinal pigment epithelium [RPE], fat, gut, ovaries/testes, heart, head kidney, liver, muscle, skin, spinal cord, spleen) were analyzed across 6 age groups spanning from a population survival of 100% (47 days) to ∼20% (162 days) (Extended Data Fig. 1a). Both males and females were sampled at a similar frequency for most tissues (Extended Data Fig. 1b), and this sex-balanced feature allowed us to study the effect of biological sex during killifish aging. Using a high-sensitivity, high-throughput library preparation pipeline based on Smart-seq2^40^, we generated a high-quality dataset, with over 94% of samples sequenced to >30 million paired-end reads and over 80% of samples having over 70% reads uniquely mapped to the killifish genome. Principal component analysis (PCA) also showed sample clustering by tissue type (Fig. 1b), confirming the tissue identity of each sample.

**Figure 1:**
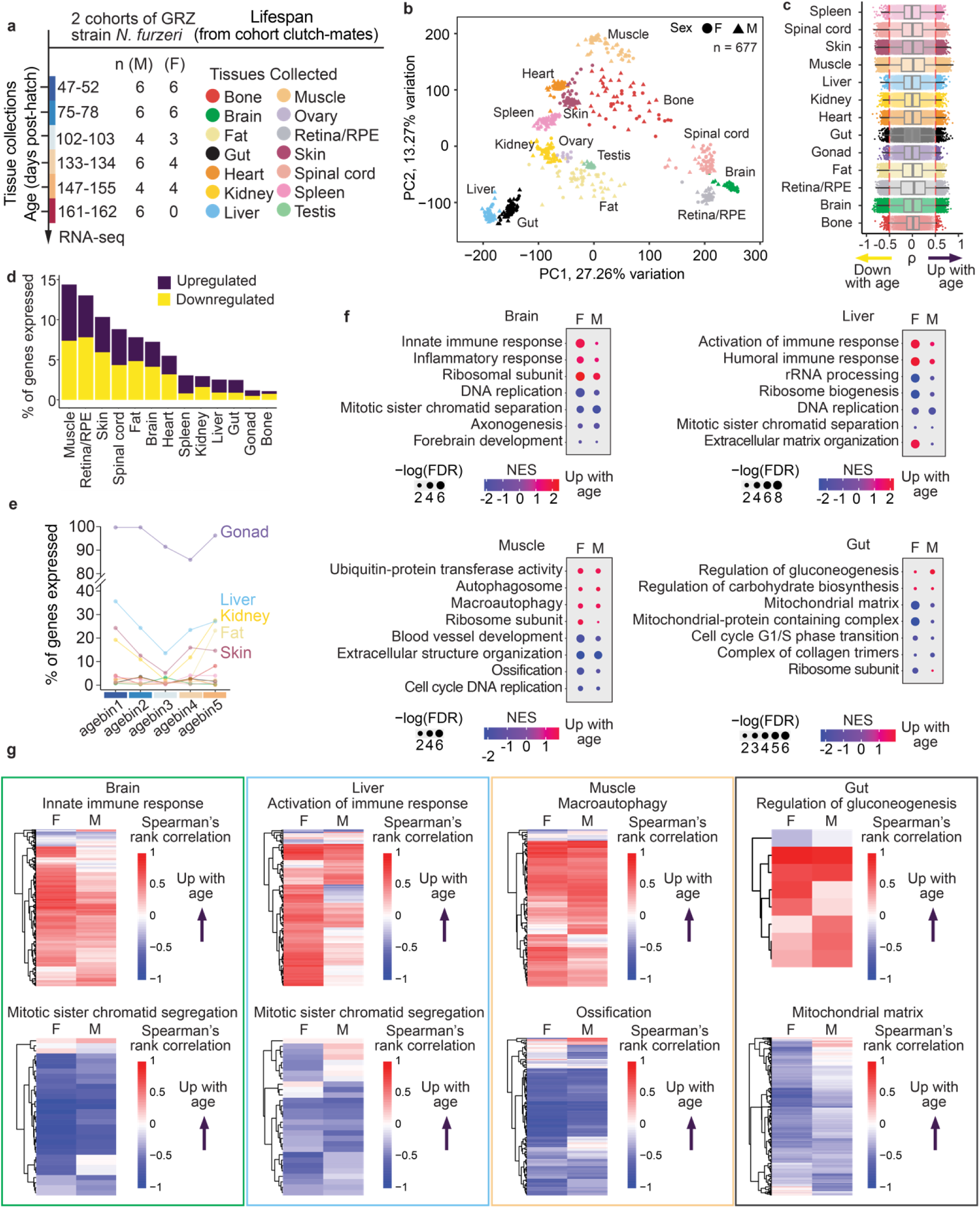
A multi-tissue killifish transcriptomic aging atlas reveals shared and tissue-specific of age effect on different tissues. (a) Schematic for the killifish transcriptomic aging atlas. Thirteen tissues from males and females were collected for RNA-sequencing at the indicated timepoints (the animal numbers sampled are listed) from two independent cohorts of the GRZ killifish strain. (b) Principal component analysis (PCA) for the 677 samples reveals clear clustering by tissue identity. Symbol shape, biological sex (F, female; M, male). Symbol color, tissue type. (c) Tissues have varying numbers of age-correlated genes, as shown by the Spearman’s rank correlation (*ρ*) distribution for all the post-filtered genes in each tissue. Each dot is one gene. Male and female samples are analyzed together for each tissue and time point in panels c to e. (d) Each tissue has distinct proportion of age-correlated genes in its transcriptome. Upregulated with age, Spearman’s rank correlation *ρ* > 0.5. Downregulated with age, *ρ* < −0.5. (e) Proportion of differentially expressed genes between males and females (sex-dimorphic genes) for each tissue, at each binned age level. A break in the y-axis is denoted by double slashed lines. (f) Male (M) vs. female (F) gene set enrichment analysis (GSEA) results, identifying the pathways significantly enriched for the genes upregulated or downregulated with age in each tissue. NES, normalized enrichment score. Dot size, –log10 of the adjusted p-value (i.e., false discovery rate [FDR] after multiple hypotheses testing). (g) Heatmap of select GO terms, plotting the male and female Spearman’s rank correlations of the genes that drive each GO term.

To characterize the gene expression trends across age in each tissue, we leveraged the time-series nature of our dataset and used Spearman’s rank correlation to describe the strength of a gene changing monotonically with age (i.e., expression consistently increasing or decreasing). Tissues such as muscle, skin, and the retina/RPE had genes with the strongest age association, with genes achieving a Spearman’s rank correlation *ρ* > 0.8 (upregulated with age) or *ρ* < −0.8 (downregulated with age) (Fig. 1c). Next, we defined age-correlated genes to have an absolute Spearman’s rank correlation greater than 0.5. We observed that among all the tissues, the muscle had the highest proportion (14.38%) of age-correlated genes in its transcriptome (Fig. 1d). Other tissues (retina/RPE, skin, spinal cord, fat, brain, heart) had an intermediate level of age-correlated genes at around 6-13%. Among the tissues with a low proportion (∼2.5%) were spleen, head kidney, liver, gut, gonad, and bone. These tissue-level differences were also observed using variance partition analysis (see Methods) (Extended Data Fig. 1c, ‘Age’), highlighting the varying degree to which aging affects the transcriptomes of different tissues.

### Age-altered pathways are mostly shared between sexes, but sex-divergent ones exist

Tissue-specific changes with age can stem from the distinctive physiology and functions of each tissue, pointing to the unique aging mechanisms in specific tissue contexts and revealing potential nodes for targeted intervention against aging in each tissue. The tissue context can depend on the biological sex of the animal from which the tissue is derived, given that the different tissue transcriptomes had varying proportion of genes differing in expression between males and females (Fig. 1e). For example, the gonads had on average ∼95% genes differentially expressed by sex across all age groups (this high degree of sex-dimorphism is expected), liver had ∼25%, skin had ∼15%, head kidney had ∼14%, and fat had ∼6% (peaking at 147-155 days of life). Consistently, variance partition analysis showed that sex accounted for a noticeable fraction of transcriptional variance in the gonad (68.70%), skin (3.55%), fat (3.49%), and head kidney (1.51%) (Extended Data Fig. 1c, ‘Sex’), and the age-sex interaction (i.e., genes changed with age differently in males vs. females) accounted for a high fraction of variance in the liver (19.97%) (Extended Data Fig. 1c, ‘Sex:age’). Prominent sex effects on tissue transcriptomes have also been observed in similar tissues in mice (e.g., gonadal adipose tissue, subcutaneous adipose tissue, liver, and kidney)^8^ and in humans (e.g., visceral and subcutaneous adipose tissue, skin)^41^.

To juxtapose male versus female differences in the aging transcriptome of each tissue, we separated our datasets by tissue and sex and then calculated the Spearman’s rank correlation for each gene, followed by Gene Set Enrichment Analysis (GSEA) to identify the pathways altered by age for each tissue and each sex (‘sex-split’ analysis). Generally in a given tissue type, we found that the significantly-enriched gene ontology (GO) terms changed with age in the same direction (either upregulated or downregulated) for both sexes, regardless of how sexually dimorphic the tissue transcriptome was (e.g., See terms for the brain, a weakly sex-dimorphic organ, and the liver, a strongly sex-dimorphic organ) (Fig. 1f and Extended Data Fig. 2). The genes underlying these pathways were mostly similar between males and females, although there were differences (e.g., the genes driving the ‘mitotic sister chromatic segregation’ term in the liver were somewhat different between the sexes) (Fig. 1g), suggesting that aging alters many pathways similarly in male and female tissues, though the exact genes altered by age can be distinct.

Interestingly, there were also GO terms showing opposite signs of upregulation or downregulation with age in the two sexes, and often the change with age was significant in only one sex (‘sex-divergent’) (Extended Data Fig. 3). Depending on the tissue type, the sex-divergent GO terms were upregulated with age in either male or female. These GO terms were related to proteostasis in the gut (e.g., ‘protein quality control for misfolded or incompletely synthesized proteins,’ ‘response to unfolded protein’); inter- and intracellular transport in the heart and spleen (e.g., ‘peptide hormone secretion,’ ‘amino acid transport,’ ‘potassium ion transport’); and the ribosome in the spinal cord (e.g., ‘ribosome biogenesis,’ ‘rRNA processing’). Two of the sex-divergent GO terms, autophagy (e.g., ‘autophagosome,’ ‘lysosomal membrane’) and myeloid cell regulation (e.g., ‘neutrophil activation,’ ‘granulocyte activation’), were present in various tissues such as fat, retina/RPE, gonad, head kidney, spinal cord, and spleen. These results indicate that while aging can alter similar pathways in male and female tissues, the direction and significance of these changes can diverge by sex, reflecting the distinct ways in which males and females age at the transcriptome level.

### Some age-altered pathways are unique to each tissue

Several pathways were altered with age in only one or a few tissues. For example, in the muscle, some age-downregulated terms were related to angiogenesis (e.g., ‘blood vessel development’) and ossification (Fig. 1f and 1g, Muscle). In the gut, metabolism-related pathways were altered with age, such as ‘regulation of gluconeogenesis’ (Fig. 1f and 1g, Gut). Even though most GO terms were consistently upregulated or downregulated with age across tissues (Extended Data Fig. 2), there were also pathways with strong tissue-dependent changes with age. For example, for both sexes, ribosome-related terms (e.g., ‘ribosome,’ ‘rRNA processing’) were upregulated with age in skin and the brain, but downregulated with age in spleen, fat, and the retina/RPE. In females, the terms related to the extracellular matrix (e.g., ‘extracellular structure organization,’ ‘extracellular matrix organization) were upregulated in the liver, fat, retina/RPE, and ovary, but downregulated in skin, muscle, and bone. How ribosome- and extracellular matrix-related processes are modulated by aging may be tuned to the different demand of ribosome activity and extracellular organization and function in different tissues.

### Immune and extracellular matrix genes change with age across multiple tissues

What pathways are commonly altered with age across multiple tissues? The shared changes could indicate systemic factors that regulate aging or shared cross-tissue consequences of the aging process. We identified several pathways that were commonly altered with age in at least 6 tissues (Extended Data Fig. 2). For both sexes, these pathways included upregulation of ‘immune response’ and downregulation of cell cycle (e.g., ‘DNA replication’) and mitochondria terms (e.g., ‘mitochondrial matrix’, ‘mitochondrial gene expression’). Specifically for male, extracellular matrix-related terms (e.g., ‘extracellular matrix organization,’ ‘extracellular structure organization) were shared across tissues (Extended Data Fig. 2). These pathways have been reported to be changed with age in a subset of killifish tissues previously^19,20,28,31,32,34,38^ and are reminiscent of key hallmarks of aging, including upregulation of ‘chronic inflammaging’ and ‘cellular senescence’ and altered ‘mitochondrial functions’ and ‘intercellular communication’^1^.

Complementarily, we analyzed male and female samples together and identified 47 age-correlated genes shared across at least 6 tissues, including 22 upregulated with age (Spearman’s rank correlation *ρ* > 0.5) and 25 downregulated genes (*ρ* < −0.5) (Fig. 2a). RNA *in situ* hybridization validated the age-altered expression of two of the top shared age-correlated genes in the gut, the tissue with the highest absolute Spearman’s rank correlation for these genes. We found that the transcript of the killifish gene *LOC107373777* (hereafter referred to as *ncRNA-3777*) (Fig. 2b, left), which is predicted to encode a long non-coding RNA of unknown function, was mostly localized to the nucleus, and its level increased with age (Fig. 2c-e). In contrast, the transcript of the *IGF2BP3* gene (killifish gene name: *LOC107383282*) (Fig. 2b, right) was both nuclear and cytoplasmic, and its level decreased with age (Fig. 2f-h). The human ortholog of the *IGF2BP3* gene encodes an RNA-binding protein that promotes insulin growth factor 2 protein (IGF2) translation^42^. Consistently, the pathways enriched for the cross-tissue age-correlated genes included immune response (upregulated) and extracellular matrix organization (downregulated) terms (Fig. 2i).

**Figure 2:**
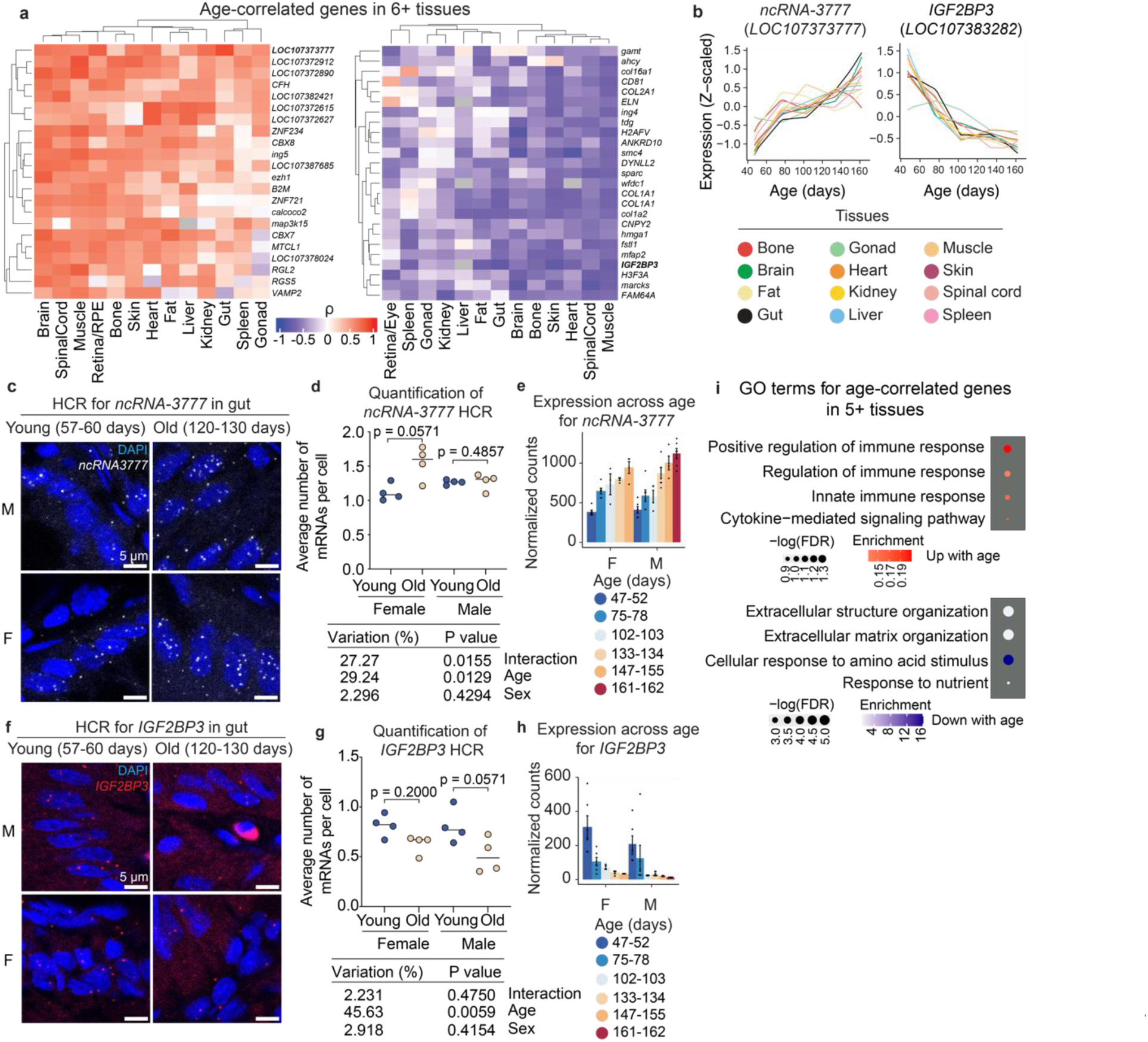
Cross-tissue comparison reveals shared age-correlated genes and pathways. (a) Spearman’s rank correlation (*ρ*) heatmaps for the genes upregulated (left) or downregulated (right) with age shared across at least 6 tissues. Gray box, Spearman’s correlation was not calculated because the expression level of a particular gene was lower than the expression threshold (TPM > 0.5 in >80% of samples). Killifish gene names are shown as lowercase letters, and additional protein-coding killifish genes are annotated using the human ortholog gene names (uppercase). The genes named after gene loci numbers (e.g., *LOC107378024*) lack human orthologs. (b) Z-scaled locally estimated scatterplot smoothing (LOESS) regression fits of the gene expression trajectories across age for the genes *ncRNA-3777* and *IGF2BP3*. (c, d) Representative maximum z-projected HCR (RNA *in situ*) images for *ncRNA-3777* and *IGF2BP3* mRNAs in male and female guts, at young (57-60 days) and old (120-130 days) ages. Scale bar, 5 µm. F, female; M, male. (e) Quantification of HCR images as the average number of *ncRNA-3777* transcripts per cell. Each dot is an animal, and four animals are analyzed for each condition. In-graph statistics, Mann-Whitney test. Below-graph statistics, two-way ANOVA with age, sex, and age-sex interaction as variables. (f) Normalized RNA-seq counts for the *ncRNA-3777* gene in the male and female guts across binned age groups. (g, h) Quantification and statistics were performed as in panels e and f, respectively, for *IGF2BP3*. Four animals are analyzed for each condition. (i) Hypergeometric GO enrichment results for the genes upregulated (top) or downregulated (bottom) with age that are shared across at least 5 tissues. Dot size, –log10 of the adjusted p-value (i.e., false discovery rate [FDR] after multiple hypotheses testing).

Next, we asked which age-altered gene pathways are conserved in mammals. Remarkably, the mouse aging atlas (Tabula Muris Senis) also reported two main categories of GO terms enriched in the top cross-tissue, age-correlated genes. The upregulated pathways were related to the immune response (e.g., ‘regulation of T cell activation,’ ‘innate immune response,’ ‘antigen processing and presentation’), and the downregulated pathways were related to intercellular interactions (e.g., ‘extracellular vesicles,’ ‘exosomes,’ ‘regulation of cell-cell adhesion’) (See figures in Schaum et al., 2020^8^, including Extended Data Fig. 2e and ‘cluster 8’ of Fig. 2a). Similarly, in a large-scale study performed in adult male cynomolgus monkeys, immune response pathways (e.g., ‘innate immune response’, ‘positive regulation of cytokine production’, ‘leukocyte mediated immunity’, ‘inflammatory response’) were also found to be upregulated with age (See ‘Cluster U’ in Yang et al., 2024^43^) and the pathway ‘extracellular matrix organization’ (See ‘Cluster D’ in Yang et al., 2024^43^) was downregulated. Additionally, an analysis of the human GTEx dataset showed upregulation of several immune pathways with age^44^. The concordance between the killifish, mouse, primate, and human data suggests that the immune system and intercellular communication (e.g. extracellular matrix, signaling) are evolutionarily conserved nodes modulated across tissues by aging in vertebrates.

### Trajectory analysis reveals different classes of gene expression behaviors

While uncovering the monotonic changes with age is informative, Spearman’s rank correlation cannot distinguish linear from nonlinear changes, nor genes with stable age trajectories from those with complex dynamics (e.g., U-shape). Previous studies revealed that age-related gene expression changes can be non-monotonic^8,26^. To explore these age-related dynamics, we performed hierarchical clustering of gene expression trajectories in each tissue, dividing the genes into 10 clusters (see Methods). We observed that the expression trajectory clusters had unique dynamics. For example, in the brain, while clusters 1, 2, and 3 all declined with age, their trajectories had distinct shapes (Fig. 3a). Cluster 1 showed a logarithmic pattern, decreasing at early age then flattening in the remaining ages. This cluster was mainly enriched in cell cycle (e.g., ‘mitotic cell cycle,’ ‘cell cycle’) and nervous system development terms (Fig. 3b, cluster 1). Cluster 2 followed a linear pattern and was enriched in pathways related to nervous system development (e.g., ‘neuron projection guidance,’ ‘neuron differentiation’) (Fig. 3b, cluster 2). Lastly, cluster 3 showed a complex behavior of declining at early age, remaining flat at middle age, and then declining further at old age. This cluster was enriched in mRNA regulation terms (e.g., ‘mRNA processing,’ ‘mRNA splicing via spliceosome’) (Fig. 3b, cluster 3). The distinct expression dynamics of these pathways may indicate different regulatory networks or the underlying reasons for the decline with age. For instance, the cluster 1 (cell cycle) pattern in the brain may result from the cessation of killifish’s rapid growth from adolescence to adulthood.

**Figure 3:**
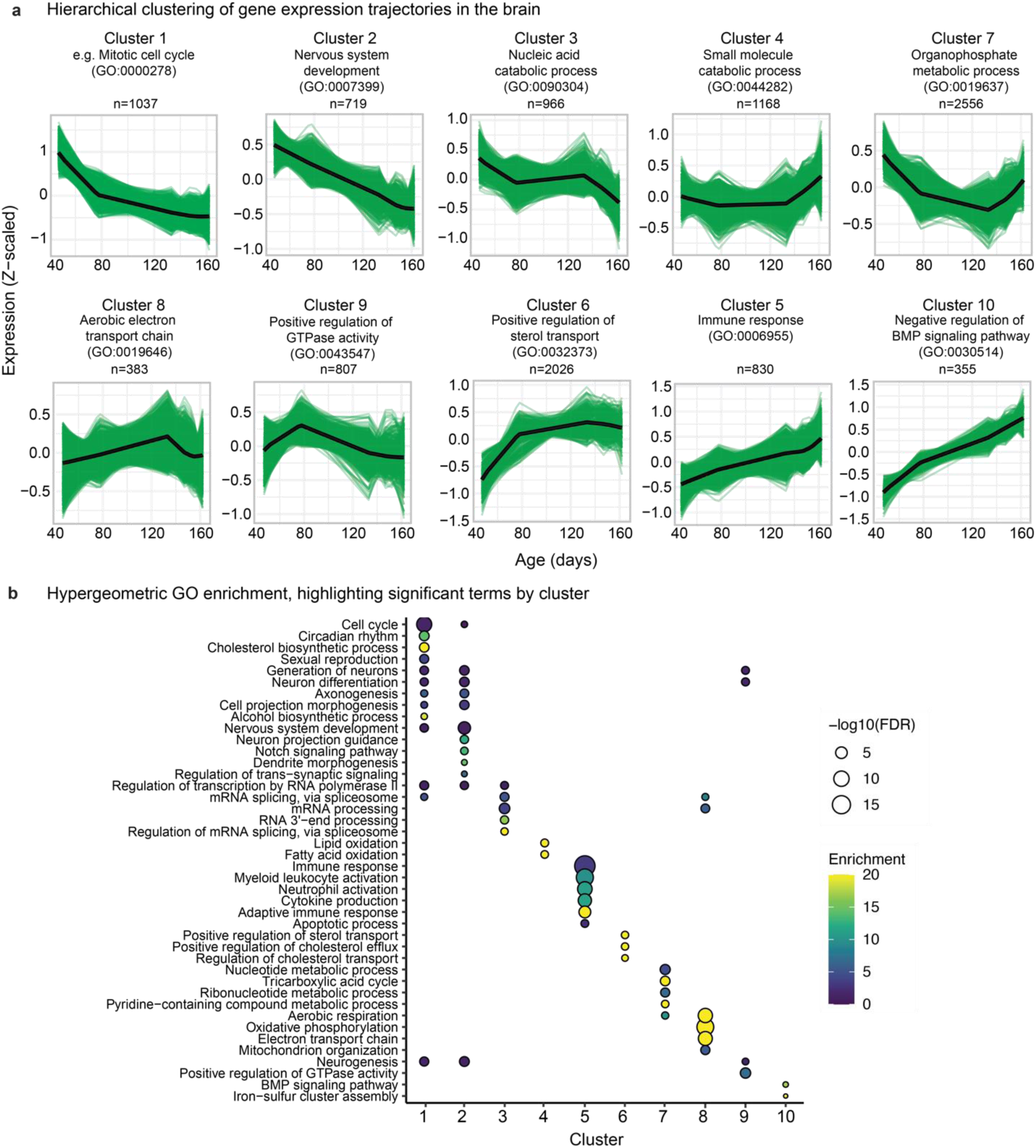
Tissue-specific gene expression dynamics for the brain. (a) Hierarchical clustering of the gene expression trajectories for the brain. Hierarchical clustering was performed on the locally estimated scatterplot smoothing (LOESS) regression aging trajectory of the gene expression in the brain for the 10,847 genes expressed in all tissues, resulting in 10 clusters of gene expression behavior over time. The average trajectory for the cluster is depicted by the black line. The most significant GO term from Hypergeometric GO enrichment (terms related to Biological Processes) for each cluster is listed. (b) Hypergeometric GO enrichment (terms related to Biological Processes) for the genes in each cluster. Select significantly enriched (adjusted p-value < 0.05) GO terms for each cluster are plotted. Dot color represents the enrichment score of each GO term, with the maximum value of the scale adjusted to 20 to improve color resolution of GO terms with lower enrichment. Dot size, –log10 of the adjusted p-value (i.e., false discovery rate [FDR] after multiple hypotheses testing). Clusters 10 does not have any significant GO terms, so the lowest p-value terms are plotted.

Consistently, other tissues had clusters with a similar logarithmic shape (an inflection point at ∼80 days) and were enriched in cell cycle pathways (e.g., cluster 8 in gut and cluster 7 in muscle) (Extended Data Fig. 4). Given that the neurogenesis terms were present in both cluster 1 and cluster 2 in the brain, it may suggest some processes related to the reduced neurogenesis as killifish age are decoupled from reduced cell division in middle-age and old brains. Lastly, the cluster 3 (mRNA regulation) pattern may reflect distinct regulatory inputs between the two phases of decline or regulation to sustain expression at middle age. Therefore, by studying gene dynamics, we can gain insights into which biological processes may be co-regulated (or not) during aging.

### Cell-type composition changes with age in the killifish kidney marrow

Given the strong systemic immune signatures, we sought to better understand how the primary hematopoietic compartment, the head kidney, of the killifish changes with age. As in other teleost fish, killifish kidneys consist of two parts. The head kidney is the anterior portion of the kidney, composed of two bilateral lobes containing hematopoietic tissue, which we sampled in our atlas. The trunk kidney is located posteriorly along the dorsal body wall and mainly contains exocrine tissue^45^. PCA analysis of the head kidney transcriptomic samples showed strong separation by age along Principal Component (PC) 1 and by sex along PC2 (Fig. 4a). We identified 516 genes with absolute Spearman’s correlation values of greater than 0.5 in the kidney samples. Several genes primarily expressed in T cells, B cells, and lymphoid progenitors were negatively correlated with age (*ρ* < 0.5), while those primarily expressed in macrophages, neutrophils, and other myeloid cells were positively correlated with age (*ρ* > 0.5) (Fig. 4b and Extended Data Fig. 5a)^46^. These differences were stronger in female head kidneys than in male head kidneys (Fig. 4b), with higher absolute Spearman’s rank correlations and greater statistical significance. At a pathway level, ‘B cell receptor signaling pathway’ and ‘DNA recombination’ terms were downregulated with age (Fig. 4c). These observations are reminiscent of the ‘myeloid bias’ phenomenon in mice and zebrafish, where the cell-type composition of the hemopoietic lineage changes with age, with an increase in the ratio of myeloid lineage cells to the lymphoid cells in old age^47–50^.

**Figure 4:**
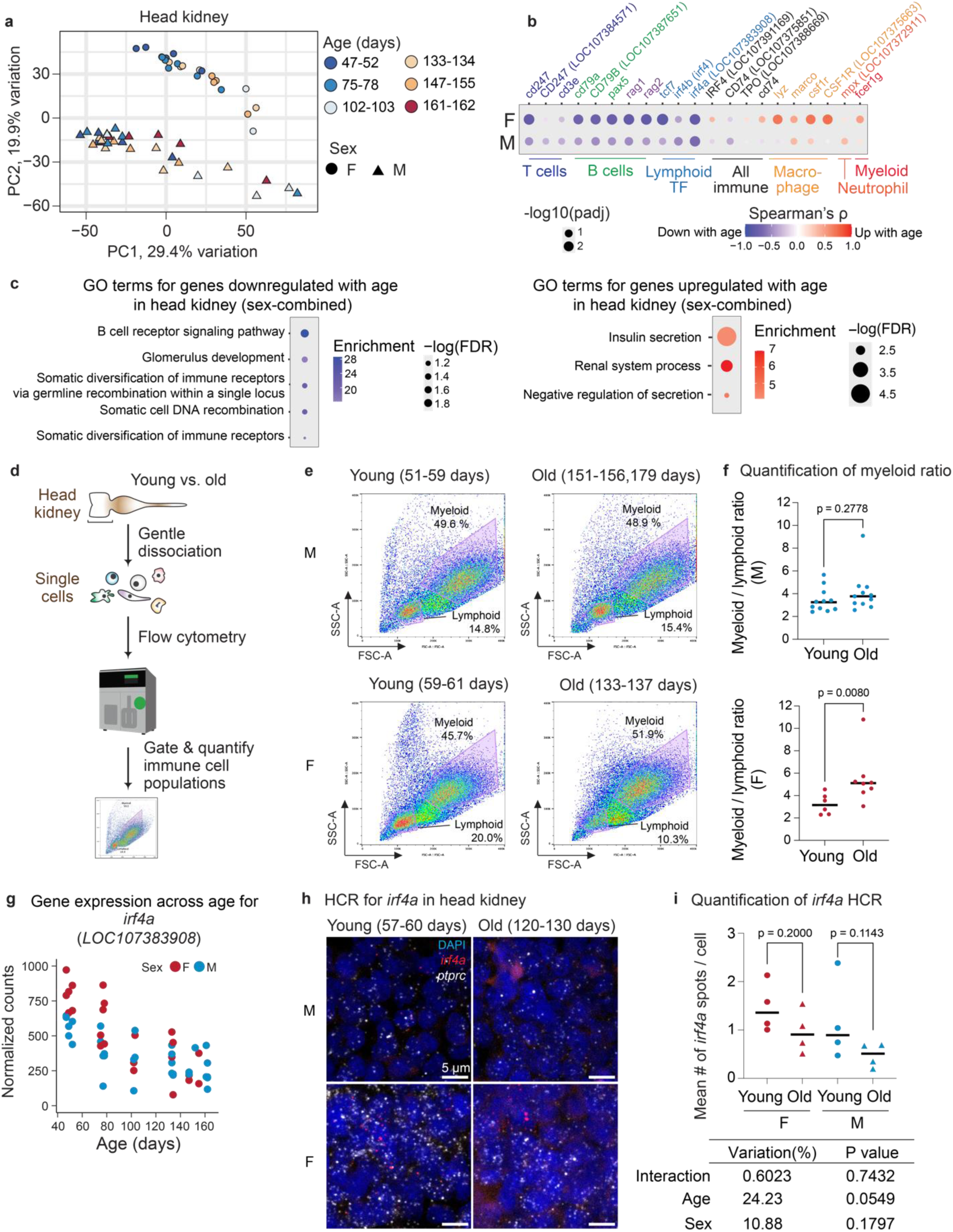
The aging killifish kidney marrow changes in gene expression and cell-type composition. (a) Principal Component (PC) Analysis of all head kidney transcriptomes coded by age (in days) and sex (female, F; male, M). (b) Dot plot of the select cell-type marker genes for lymphoid and myeloid lineage cells. If a gene is named after a gene locus number (e.g., ‘*LOC107384571*’), either the zebrafish homolog (all lowercase) or human homolog (all uppercase) is also written. The dot size is the –log10 of the adjusted p-value, and the dot color corresponds to the Spearman’s rank correlation *ρ* value calculated separately for each sex. The cell-type specificity of each gene’s expression was based on a published killifish kidney single-cell RNA-seq dataset^46^ (see Extended Data Fig. 5). (c) Hypergeometric GO enrichment (terms related to Biological Processes) for the genes upregulated (right) or downregulated (left) with age identified for the head kidney when both sexes were analyzed together. Dot color represents the enrichment score of each GO term. Dot size, –log10 of the adjusted p-value (i.e., false discovery rate [FDR] after multiple hypotheses testing). (d) Schematic of the flow cytometry assay to quantify different immune cell lineages in the killifish. Dissected head kidney tissue was dissociated into a single-cell suspension and analyzed by Fluorescence Activated Cell Sorting (FACS). (e) Representative forward-scatter vs side-scatter flow cytometry plots from male and female killifish. Myeloid and lymphoid gates are depicted as the percentage of total live cells. (f) Quantification of myeloid: lymphoid ratio (total myeloid events: total lymphoid events) from flow cytometry data. Each dot is an animal, and 12 males and 6-8 females at each time point were analyzed for panels e and f. Significance determined by Mann-Whitney test. (g) Scatterplot of the counts normalized by DESeq2 for *irf4a* (*LOC107383908*), with each dot representing the expression of *irf4a* in an individual sample in the atlas dataset. Red, female (F). Blue, male (M). (h) Representative maximum z-projected HCR images of male (top) and female (bottom) kidney sections at young or old ages. The sections were stained with DAPI (blue) and the HCR probes against *irf4a* (red) and *ptprc* (white) mRNAs. Scale bars, 5 µm. (i) Quantification of the HCR images in panel h. The average number of *irf4a* mRNAs per cell is plotted (only the interstitial regions were quantified). Each dot is an animal (4 animal per each sex and age group were quantified). In-graph statistics, Mann-Whitney test. Below-graph statistics, two-way ANOVA with age, sex, and age-sex interaction as variables.

To test whether the changes in the killifish head kidney gene expression were due to cell-type compositional changes (e.g., ‘myeloid bias’), we optimized a head kidney dissociation protocol followed by fluorescence activated cell sorting (FACS) (Fig. 4d and Extended Data Fig. 5b). We validated a FACS gating strategy developed for zebrafish (based on forward- and side-scatter^51^) by performing RNA-sequencing on the FACS-sorted cells and found enrichment for either lymphoid or myeloid cell-type specific expression in the expected cell populations (Extended Data Fig. 5c and 5d). Using this strategy, we observed that females, but not males, exhibited age-related cell-type compositional changes (Fig. 4e and 4f). There was a significant increase in the ratio of putative myeloid to putative lymphoid cells in old females (133-137 days old) compared to young females (59-61 days old) (p = 0.0080), whereas such increase was subtle and not significant in males (151-179 days vs. 51-59 days of age) (p = 0.2778). This more pronounced cell-type compositional change in females is consistent with the stronger age correlation observed in gene expression for females (Fig. 4b). Such sex differences may occur because the females in our cohorts were shorter-lived than males (Extended Data Fig. 1a) and likely aged more rapidly than males. Interestingly, among the most strongly downregulated genes were the two orthologs of the lymphoid transcription factor *IRF4* gene in mammals^52–54^ and zebrafish^55^ (Fig. 4b). These two killifish paralogs of *IRF4*, *irf4a* (killifish name: *LOC107383908*) and *irf4b* (killifish name: *irf4*), have differing expression levels and patterns (Fig. 4g, Extended Data Fig. 5e and 5f)^46^, with *irf4a* more strongly downregulated with age (Fig. 4b). We validated the *irf4a* transcript levels by RNA *in situ* hybridization, showing that *irf4a* mRNA could be co-expressed with *ptprc* mRNA (*CD45,* a pan-leukocyte marker) in cells of the hematopoietic-tissue-enriched interstitial regions of the killifish head kidney (Extended Data Fig. 5g and 5h) and decreased with age (two-way ANOVA, p = 0.0549 for the ‘age’ variable) (Fig. 4h and 4i). While we could not validate Irf4a protein expression (no fish-specific Irf4a antibody exists currently), our results raise an interesting possibility that *irf4a* downregulation with age may reduce lymphoid cell differentiation, leading to increased relative abundance of myeloid cells.

### Biological age can be predicted using tissue-specific clocks

Our comprehensive transcriptomic aging atlas allows us to develop age-prediction models for each tissue, known as ‘aging clocks’^56–59^. Using molecular features from large datasets (e.g., DNA methylation^43,60–62^, transcriptomes^43,63,64^, proteomes^65^), these machine-learning models first learn patterns from samples of known chronological ages (‘training’) and then compare the molecular pattern of a query sample (which is not used in the training set) with the learned patterns to find the age best matched by the query, the ‘predicted age.’ Development of these clocks has accelerated evaluation of genetic, pharmacological, and lifestyle aging interventions. For example, the epigenetic aging clocks trained on chronological age predict animals and humans to have ‘younger’ age when they are subjected to beneficial health interventions such as diet and exercise^61,66–68^ and lifespan-extending genetic manipulations^62,69,70^.

To build tissue-specific transcriptomic aging clocks, we used three machine-learning modeling strategies, including the Bayesian, non-linear pipeline BayesAge 2.0^71^ (Fig. 5a), Elastic Net regression (a hybrid model of LASSO and Ridge regression) (Extended Data Fig. 6a), and Principal component-based regression^72^ (PC-R, Extended Data Fig. 6b) (see Methods). Applied to our dataset, these models had different prediction precision and residual behaviors (whether a model’s predictions underestimate or overestimate the true values) (Extended Data Fig. 6c-g), and thus we reported the results of all three. For example, for BayesAge 2.0 and Elastic Net, the gut and testis were among the highest performing clocks, with correlation coefficients (R^2^) over 0.8 (Fig. 5b, Extended Data Fig. 6a and 6b). The lowest performing clock was the ovary clock, likely because our dataset has fewer samples for the majority of timepoints for this tissue, due to sample dropout (Extended Data Fig. 1b, 6h-i and Methods).

**Figure 5:**
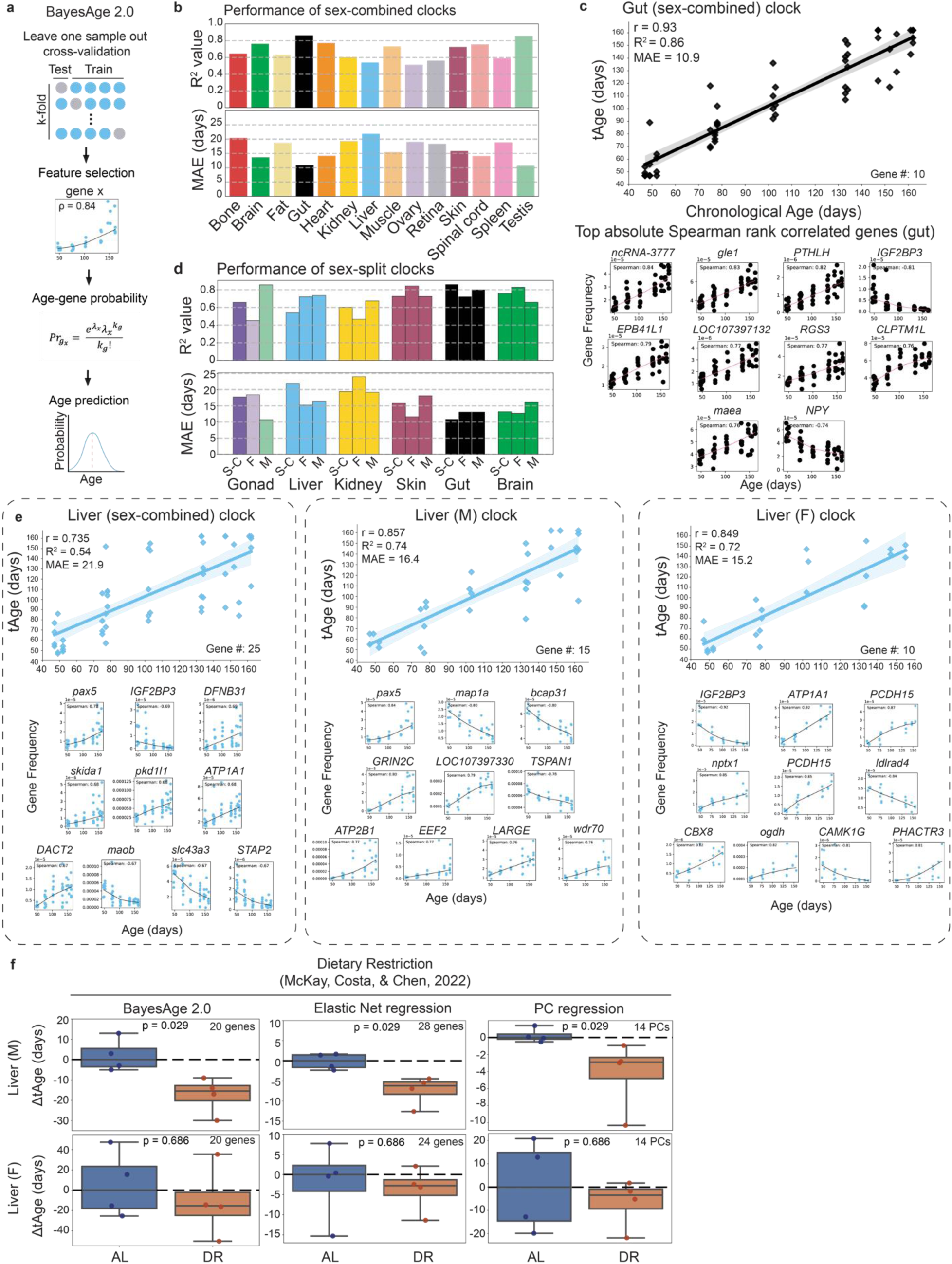
Tissue-specific transcriptomic aging clocks predicts tissue biological ages in aging and interventions. (a) Workflow of BayesAge 2.0, a Bayesian and locally weighted scatterplot smoothing (LOWESS) regression model behind the aging clocks. To train a tissue clock, Leave One Sample Out Cross-Validation (LOSO-CV) was used to generate testing-training splits of the data. In each iteration of LOSO-CV, one sample was used as a test set, while the rest of the tissue samples were used for training. This was performed k times, where k is the number of tissue samples available. Each time LOSO-CV was performed, a set of top age-associated genes (the highest absolute Spearman’s rank correlation values) was selected for the feature set. Then, the probability that the sample in the test set was a given age was calculated from the probability of the observed expression value for each selected gene in the sample at that age, assuming a Poisson distribution. The product of each gene-wise probability was computed to determine the age probability. The result was an age-probability distribution from which the age prediction was the highest probability age in this distribution. (b) Bar plots of the performance metrics for the BayesAge sex-combined tissue clocks, using the coefficient of determination (R^2^) for the relationship between chronological and predicted age and the mean absolute error (MAE). (c) Scatterplot of gut clock chronological age vs. the ‘transcriptomic age’ (tAge) for measuring the prediction accuracy of the highest performing gut sex-combined tissue clock. The ‘optimal’ BayesAge clock is defined as the model with the most concordance between chronological and predicted age among all the gene number tested. Bottom, the gene frequency scatterplots of the top 10 overall age-correlated genes trained on the sex-combined gut samples are shown. The pink line is the locally estimated scatterplot smoothing (LOESS) regression fit across time. (d) Bar plots of R^2^ and MAE values for select clocks trained on sex-combined data (left, ‘S-C’), female data (middle, ‘F’), and male data (right, ‘M’). Selected tissues include highly transcriptionally sex-dimorphic tissues (gonad, kidney, liver), moderately transcriptionally sex-dimorphic tissues (gut, skin), and one weakly sex-dimorphic tissue (brain). (e) Accuracy of tAge predictions for the optimal sex-combined (left), male-only (middle), and female-only liver clocks (right). (f) Predicted ages for liver samples from male and female killifish fed on *ad libitum* (AL) or dietary restricted (DR) diets using sex-dimorphic liver clocks (data from a published dataset^22^). Age prediction was performed using three different modeling strategies, BayesAge 2.0 (left), Elastic Net regression (middle), and Principal Component regression (right). Each dot in each box plot represents the predicted tAge for the liver transcriptome of an individual fish (4 fish per condition) and the gene set size or number of principal components used for age prediction is listed. For each model, Mann-Whitney test was used to test the significance of difference between the AL and DR conditions.

What age-correlated genes are driving the aging clock of each tissue? We examined the top 10 genes underlying the aging clocks for some of the top performing BayesAge 2.0 models (gut, brain, and testis) (Fig. 5c and Extended Data Fig. 7a-d). Generally, the human orthologs of these genes were functionally related, possibly reflecting key functional changes in aging. For example, the top 10 gut genes were related to nutrient sensing, including neuroendocrine peptides *PTHLH* and *NPY* and the *IGF2BP3* gene, which encodes an IGF2 translation regulator protein^42^ (Fig. 5c, bottom). For the brain, several of the top genes have been reported to regulate cell division, such as *CENPF*, *SMC4*, and *RCC2* (Extended Data Fig. 7a and 7b), and *DLL1* has been implicated in adult neural stem cell maintenance^73^. These genes are consistent with the reduced neurogenic capacity of the aged killifish brain, as reported previously^35^. Finally, the top testis genes were related to cytoskeleton functions, including *GSN*, *KRT8*, and two orthologs for *TUBB4B* (Extended Data Fig. 7c and 7d). Together, we find that for the best performing tissue clocks, the genes underlying the clocks share related functions, hinting at key regulators of tissue-specific aging dynamics.

Because our dataset is relatively sex-balanced, for each tissue, we compared the performance of the aging clocks developed using each sex’s transcriptome (‘sex-split’) with those built from sex-combined transcriptomes. Interestingly, for the liver, both sex-split BayesAge 2.0 clocks outperformed the sex-combined clock, improving the R^2^ values from 0.735 (sex-combined) to 0.857 (males) and 0.849 (females) (Fig. 5d and 5e). For other sex-dimorphic tissues (e.g., head kidney, skin, and gonads), the BayesAge sex-split clocks improved the clock performance of only one sex (male or female) (Fig. 5d). This improvement could also occur for a less sex-dimorphic tissue, such as the brain (Fig. 5d). Therefore, while sex-split clocks do not always improve the clock performance of sex-dimorphic tissues, they can in specific cases (e.g., liver). In addition to sex-combined models, sex-split models should be tested for developing better age-prediction models.

Lastly, to test the utility of our transcriptomic clocks, we used the clocks to make age predictions on a published transcriptomic dataset of a lifespan-extending intervention. Previously, we reported a dietary restriction paradigm (‘DR’) that extends male lifespan in killifish by 16-22% but has no effect on female lifespan^22^. Our sex-split liver clocks (using all three machine-learning modeling strategies) revealed that for males, DR significantly decreased the predicted age of the liver sample transcriptomes (ΔtAge) in comparison to the *ad libitum* (‘AL’) paradigm (p = 0.029 by BayesAge 2.0, p = 0.029 by EN, and p = 0.029 by PC-R) (Fig. 5f and Extended Data Fig. 8a-d). In contrast, for females, DR did not significantly decrease the predicted age of the liver transcriptome (ΔtAge) in comparison to AL (p = 0.686 by BayesAge 2.0, p = 0.686 by EN, and p = 0.686 by PC-R) (Fig. 5f and Extended Data Fig. 8a-d). This finding is consistent with the observation that this DR paradigm does not extend female lifespan^22^. Therefore, the transcriptomic clocks can make age predictions on unseen data, consistent with biological contexts and providing insights into biological age.

## Discussion

We have presented a comprehensive aging transcriptome atlas of 13 tissues for male and female killifish. To facilitate sharing of this useful resource, we have compiled all the results on an open-access online portal (see Methods). Our analyses reveal varying age-sex relationships for each tissue, identifying several sex-dimorphic tissues (e.g., gonads, liver, gut, head kidney) that benefit from analyzing each sex separately. Time-series correlation analysis and gene expression trajectory analysis have identified age-correlated genes and pathways shared across multiple tissues, including several ‘hallmarks of aging’ related to inflammation, extracellular matrix, mitochondria, and proteostasis. Importantly, these hallmarks are consistent with the findings in mammals, such as those reported in the mouse aging atlas Tabula Muris Senis^8^, suggesting evolutionary conservation between killifish and mammals.

In our study, most of the age-altered pathways are consistent between males and females. One of the strongest pathways upregulated with age is related to immune response. Both innate and adaptive immune responses are elevated in old males and females across at least six tissues. This upregulation may be driven, in part, by increased immune cell infiltration, which is reported in several killifish tissues^20,35,36^. Recently, single-cell datasets have become available for several killifish tissues^21,35,46,74^, including the kidney. Integration of our bulk RNA-sequencing data with these single-cell data using computational deconvolution techniques^75,76^ can help distinguish shifts in cell-type composition from gene expression changes in each cell type. Furthermore, it is possible that the level of cross-tissue inflammation elevation may be linked to the degree of cell-type composition changes in the hematopoietic tissue (head kidney). For instance, females have a stronger increase in the relative proportion of myeloid cells with age compared to males, and correspondingly, more tissues upregulate innate immune responses in females than in males. It will be interesting to further explore what explains the gene expression and cell-type composition changes with age in the head kidney and how altering kidney aging may influence systemic inflammation of other tissues.

Another interesting class of age-altered pathways is related to the extracellular matrix (ECM), which are downregulated with age in almost all the tissues in males and in a subset of tissues in females. The ECM plays a central role in tissue structural maintenance and cell-cell signaling and is impacted by aging in animals and humans. For example, ECM genes (transcripts and proteins) are altered with age in mice, primates, and humans^8,43,77,78^. While ECM disruption can accelerate aging in mice^79,80^, longevity interventions have been shown to promote ECM homeostasis in *C. elegans*^81^. While lifespan extension has not been shown by modulating the ECM in vertebrates, our study, along with others in the literature, highlight the growing body of evidence for a role of the ECM in regulating aging in animals.

In addition to the aging pathways shared between males and females, there are also pathways that diverge in their directions of change (upregulated or downregulated with age) between the two sexes. Interestingly, male and female killifish often differ in their responses to lifespan interventions (so that lifespan is extended in only one sex), including dietary restriction and intermittent fasting^19,22^, genetic mutations in the AMPK pathway^19,20^ and the germline^21^, and metformin treatment^27^. The sex-divergent pathways may contribute to the sex-specific responses to lifespan interventions. It would be interesting to screen interventions in a sex-specific manner (e.g., testing small molecules that specifically target female pathways). Excitingly, our tissue-specific transcriptomic aging clocks, which include sex-split models, can accelerate evaluation of the efficacy of interventions by using transcriptomic signatures as a readout (instead of lifespan). It will be useful to apply our transcriptomic tissue clocks on additional datasets that involve genetic mutants to test how broadly these clocks can capture different aging interventions. We envision that this comprehensive transcriptomic atlas and the associated aging clocks will not only accelerate discovery of drivers and biomarkers of tissue aging but also enable the rapid evaluation of future aging interventions in the killifish, an powerful short-lived vertebrate model for aging research. Furthermore, these resources should help identify shared aging pathways across species.

## Contributions

E.K.C., J.C., I.H.G., A.B., and T.W.-C. conceptualized of the study. E.K.C., J.C., and I.H.G. raised animals for cohorts, designed collection strategy, and harvested all atlas tissues. I.H.G. optimized the transcardial perfusion protocol and perfused each animal in this study. E.K.C., J.C., and I.H.G. performed RNA extractions, E.K.C. and J.C. performed library preparations and all the computational analysis except the tissue clocks. L.M. designed the computational method BayesAge 2.0 under the supervision of M.P. and L.S.B. and worked with E.K.C. to refine the clocks. E.K.C. curated independent query datasets for implementation of age prediction using tissue clocks. J.C. performed validation experiments and collected tissue samples. E.K.C. and N.S. performed histological sectioning. E.K.C., N.S., A.T., J.C. performed HCR staining, imaging, and analysis. M.R.W. performed retina/RPE dissections and RNA extractions under the supervision of S.W. E.K.C. and J.C. performed tissue dissection and FACS experiments on head kidney. P.M.S. made the Shiny App for data exploration and advised on data preprocessing. P.P.S. provided the general RNA-sequencing analysis pipeline (quality control, mapping, DESeq2, GSEA analysis) and provided computational analysis advice. E.K.C., J.C., A.B., and T.W.-C. wrote the original manuscript draft. A.B. and T.W.-C. supervised the study.

## Data Availability

The raw FASTQ files will become public in the Sequence Read Archive (SRA) upon publication. The normalized expression data matrix is available under the same SRA accession and for exploration through a R-based Shiny application: https://twc-stanford.shinyapps.io/atlas/. Raw images will be deposited to figshare and will become public upon publication.

## Code Availability

All code has been shared in the public GitHub repository https://github.com/emkcosta/KillifishAtlas.

## Supporting information

Submission_Files_250120

## Acknowledgements

We thank all members of the Wyss-Coray and Brunet Labs, particularly Felix Boos, Rahul Nagvekar, Nimrod Rappoport, Eric Sun, and Andy Tsai for scientific discussion and feedback on the manuscript. We thank Felix Boos, Nimrod Rappoport, and Patricia Moran-Losada for help with independent code validation. We thank Justin You for discussion on image quantification parameters. We also thank Maddie Housh, S.V. Perry, Sarah Boyle, Jacob Chung, Rishad Khondker, and Rogelio Barajas, for their assistance with killifish husbandry over the years. We thank Divya Channappa, Kathleen Dickey, and Hui Zhang for administrative support. This worked was supported by The Phil and Penny Knight Initiative for Brain Resilience (T.W.-C.), the NOMIS Foundation (T.W.-C. and A.B.), the Milky Way Research Foundation (A.B.), the Glenn Foundation for Medical Research (A.B.), and the CZ Biohub Investigator Program (A.B.). E.K.C. was supported by the National Institutes of Health (NIH) Training Grant (T32MH020016) awarded to the Leland Stanford Junior University and the Fondation Bertarelli Graduate Fellowship Fund. J.C. is a Jane Coffin Childs fellow, Stanford Katharine McCormick fellow, and a Stanford Jump Start Award recipient. I.H.G. is funded by an NIH Pathway to Independence Award (1K99AG088304-01) and a MAC3 Dementia and Ageing Fellowship. L.M. is a Eugene V. Cota-Robles fellow, Warren Alpert Computational Biology and Artificial Intelligence Network scholar, and National Science Foundation (NSF) Research Traineeship (NRT) fellow. This work was additionally supported by the NIH Training Grant in Genomic Analysis and Interpretation (T32HG002536) (L.M.), and the National Science Foundation (NSF) UCLA Quantum Science and Engineering PhD Fellowship (L.M.).

## Additional Information

Competing interests: No competing interests.

## Materials and Methods

### African turquoise killifish husbandry

All experiments used the GRZ strain of the African turquoise killifish species *Nothobranchius furzeri*. Fish were housed in a 26°C circulating water system kept at a conductivity between 3500 and 4500 μS/cm and a pH between 6.5 and 7.5, with a daily water exchange of 10% with reverse-osmosis-treated water. All animals were kept on a 12 h/12 h day/night cycle. Feeding and husbandry details are described below. All fish were housed within the Stanford Research Animal Facility under protocols approved by the Stanford Administrative Panel on Laboratory Animal Care (IACUC protocols #31727 and #13645).

#### Atlas cohorts

All fish were raised from embryos collected from group breeding tanks (1 male paired with at least 3 females in 9.8 L tanks, and the breeders are generally 2-4 months old). Breeder tanks were fed ∼18 mg Otohime fish pellets per fish (Reed Mariculture, Otohime C1) twice a day and bred with sand trays in the tanks for embryo collection. After 4-8 h, the sand trays were collected, and embryos were separated from the sand by sieving. To reduce contamination, we rinsed the embryos with 0.2% mild iodine (diluted from Povidone-iodine solution [10% w/v, 1% w/v available iodine, RICCA 3955–16] in Ringer’s solution [Sigma-Aldrich, 96724]). Decontaminated embryos were incubated in Ringer’s solution supplemented with 0.01% methylene blue (Kordo, 37344) at 28°C in 60 mm x 15 mm Petri dishes (E and K Scientific, EK-36161) at a density between 10 and 50 embryos per plate for ∼2 weeks and then placed on moist coconut fiber substrate (Amazon, B00167VVP4) at 26°C. After ∼2 weeks on coconut fiber, fish were hatched in ∼1 cm-deep chilled (4°C) 1 g/L humic acid solution (Sigma-Aldrich, 53680) and incubated at room temperature overnight. For the next 4 days, the hatched fish were housed at room temperature. During this period, system water was added to the hatching containers, and fish were fed 2-3 drops of live brine shrimp (hatched from Premium grade brine shrimp eggs [Brine Shrimp direct, 12-pound carton], see published protocols for details^82^) once daily using plastic pipettes (Globe scientific, 138090). Fish were housed at a density of 4 fish per 0.8 L tank for the following two weeks, then 2 fish per 0.8 L tank for one week, and then 1 fish per 0.8 L tank for one week. Fish were fed with brine shrimp twice daily. At the 5th week post-hatching, each fish was transferred to a 2.8 L tank and sexed by caudal fin color: males exhibit vivid colors, but females do not. Fish with severe gill defects, curved spines, and an inability to float (‘belly sliders’) were excluded. A random subset of individuals from each cohort were designated as ‘Lifespan’ animals, and these animals were not selected for harvest. Any other unharvested animals that died from natural causes were also plotted in the lifespan analysis. Cohort 1 fish were enrolled in two batches, 2 weeks apart (See Supplemental File 1 for enrollment details). Cohort 2 fish were enrolled as an independent cohort, 6 months apart from Cohort 1. All fish from each cohort were randomly assigned to tank locations using the ‘Randomize Range’ function in Google Sheets. Cohort 1 (first enrollment) fish were fed using an automated feeder^22^ under the *ad libitum* regimen (5 mg per feeding and fed 7 times a day for a total of 35 mg of Otohime fish pellets). Cohort 1 (second enrollment) and Cohort 2 were fed using a custom-made manual feeder twice a day, 18 mg per feeding, for a total of 36 mg of Otohime fish pellets. The core design of the custom feeder has the same acrylic-cut feeding disc as the automated feeder, and thus, it has the same precision as the automated feeder.

#### Validation cohort for RNA in situ staining

Fish were raised similarly to the atlas cohorts with the following modifications. After collection, embryos were rinsed several times with embryo solution (Ringer’s solution with 0.01% methylene blue) instead of mild iodine, placed in fresh embryo solution, and incubated at 26-28°C. Approximately two weeks after collection, embryos were placed on moist coconut fiber and incubated at 27°C. Two weeks later, fish were hatched in 60 x 15 mm Petri dishes (VWR, 25384-168) containing 10 mL of cold 1 g/L humic acid solution and placed at room temperature on the bench top. After the fish were hatched, they were placed into the 26°C circulating water systems in 0.8 L tanks at a density of 10-20 fish and fed brine shrimp twice daily. After one week, the fish were split and housed at a density of 4 fish per 0.8 L tank for one week, then 2 fish per 0.8 L tank for another week, and then 1 fish per 0.8 L tank for one week. At the 5th week post-hatching, fish were upgraded to 2.8 L tanks, sexed, and randomly assigned to their tank positions. All sexually mature fish were fed using the custom-made manual feeder as in the atlas cohorts (18 mg of dry pellets twice a day, for a total of 36 mg per day). We note that the validation cohort was run as the control for another experiment, which aimed to understand how mating affects killifish aging, and the validation fish were the ‘unmated control.’ Thus, the validation animals were housed with sand trays (which were used as the mating bedding for the mated group) for 4 h twice a week (8 h total per week). The male fish experienced a ‘mock cross’ twice a week, where the male fish were netted and placed back to their own tanks to mimic the ‘crossing’ of the mated group.

### Lifespan analysis, including Kaplan Meier curve plotting

First, animals with missing data (e.g., for sex or death date) or those harvested for RNA-sequencing were excluded from the analysis. The remaining animals (the animals designated for lifespan analysis and those that died of natural causes) were used to plot Kaplan Meier survival curves. Data was entered into Prism using the defaults for survival analysis, with ‘1’ being used for a censored sample and ‘0’ for when a sample died. Kaplan Meier curves were plotted individually for males and females, separated by enrollment Cohort.

### Atlas cohort tissue collection

The harvest dates were randomly assigned to each fish within each cohort. On a harvest day, each fish was fed 18 mg of Otohime fish pellets 7:30 - 8 AM. At ∼10:30 AM, the fish were transported from the animal facility to the lab space in their own tanks. Typically, 4 fish (2 males and 2 females) were dissected on each harvest day (∼30 min to dissect each fish). Dissection began with perfusion (see details below) and then tissue collections on top of ice-cold Sylgard-coated Petri dishes (filled with wet ice and covered in plastic wrap) by three operators (E.K.C., J.C., I.H.G.). Dissected tissues were placed in 1.5 mL tubes (Fisherbrand, 02-681-320), snap-frozen in liquid nitrogen, and stored at -80°C until RNA extraction. The same operator dissected the same tissues for all fish in this study (See Supplemental File 1 for details). Muscle samples were collected from a ∼1 cm region immediately anterior to the caudal fin, with skin removed and cuts made above and below the horizontal septum to remove the spinal cord and vertebrae. The spinal cord was collected by dissecting out the vertebrae and gently pulling the spinal cord from the vertebral foramen. Skin samples corresponded to the caudal fin’s most posterior ∼0.5 cm portion. The retina and retinal pigment epithelium (RPE) were dissected from the eye together (by M.R.W.). In some cases, the retina/RPE samples were dissected from individual eyes from the same animal, and in other cases, samples were pooled between animals (indicated in Supplemental File 2 where relevant). Only the head kidney was collected for the kidney samples. For the liver samples, the pale green gallbladder was removed whenever it was visible. Total visceral fat was collected (without regional distinction). All oocytes were collected for ovaries, including those that had fallen out of the organ during dissection.

#### Perfusion Device Setup

To perfuse a killifish, a syringe pump (KD Scientific, Legato 200 Series, 788200) that permits hands-free depression of the perfusion syringe was set up as follows: a 20-mL disposable syringe with Luer Lock tip (‘Sterile Syringe Only with Luer Lock Tip’, Amazon, B08FJCSLFC) was attached to a 30 gauge metal hub blunt-end Luer needle (Hamilton Syringe, custom needle, 7748-16; 30 gauge, Metal Hub Needle, Point Style: 3; Needle Length: 0.375 inches). The blunt-end Luer needle was connected to ∼0.25 meters of BD Intramedic PE Tubing (BD, 427400), which terminated in a 30-gauge hubless needle with a point style 4 bevel (Hamilton Syringe, custom needle, 22030-01; 30 gauge, Hubless Needle, No Hub, 30 gauge, 1.5 inches length, point style 4 [12°]). The 20 mL syringe was filled with nuclease-free 0.25 M EDTA diluted in 1x PBS (Corning, 21-040-CV) and fitted into the syringe pump.

#### Killifish Perfusion

The killifish was first deeply anesthetized in tricaine (100 mg/L system water, pH ∼7 using sodium bicarbonate) until operculum movement slowed, and the fish was unresponsive to touch. Once deeply anesthetized, the fish was placed on top of a Sylgard-coated Petri dish filled with wet ice covered in plastic wrap. The fish was secured on its side with two dissection pins – one pin piercing the muscle immediately anterior to the caudal fin and one pin piercing the gill operculum that lay in contact with the plastic wrap.

First, the gill was exposed by cutting off the operculum with scissors. Operculum removal helps visualize the gill and evaluate perfusion completion, as the gill would be flushed of blood and turn white with successful perfusion. Next, using a scalpel, a small ∼1 mm incision was made through the skin immediately anterior to the urogenital opening. A scissor was then inserted at the incision site and cut along the ventral side of the fish to the gill, only cutting through the skin. Next, using the ventral incision as a starting point, a ‘window’ was created using scissors to remove the body wall covering the liver and heart. Once the heart was visible, Iris forceps were used to gently remove the transparent membrane that partially covers the heart and connected the heart to the body wall. Removal of this membrane exposed the heart for complete visibility during perfusion. Next, spring scissors were used to cut the atrium to create a blood flow outlet. Immediately after cutting the atrium, the hubless needle of the perfusion device was inserted ∼1 mm into the apex of the ventricle (or as deep as the bulbus arteriosus), and the syringe pump was switched on to depress the plunger of the syringe at a rate of 3.5 mL/minute to initiate perfusion. The needle was steadily held in place until the gill and liver were visibly perfused of blood.

### RNA isolation

To reduce within-tissue batch effects, we processed all samples of the same tissue type on the same day unless otherwise noted. Due to the large number of samples, RNA extraction was performed in 2-3 batches for each tissue, with the order of samples randomized and a roughly equal assignment of age and sex combinations to each batch. The processing order of each sample within a tissue type was randomized using the “Randomize Range” option in Google Sheets. After randomization, tissue samples were assigned unique numerical “RNA_IDs” and split into batches of 12-24 samples for processing.

The RNA isolation protocol was based on the RNeasy Mini RNA extraction protocol from QIAGEN and largely kept consistent between tissues, except for when stated otherwise below. The general RNA extraction protocol is as follows. First, tissue sample tubes were removed from -80°C storage, placed in liquid nitrogen, and transferred to a 4°C cold room to prevent tissue thawing. Sample tubes were placed on a pre-chilled (-20°C) TissueLyser 2 mL tube adapter (QIAGEN, 69982) on dry ice in the cold room, and ∼100 µL of pre-chilled at 4°C Zirconia/Silica beads (0.5 mm diameter BioSpec Products, 11079105z) were added to each tube. Next, the sample tubes were quickly transferred to wet ice on the adapter, and 700 µL of 4°C QIAzol lysis reagent (QIAGEN, 79306) was added to each tube. The sample tubes were placed between the pre-chilled (4°C) metal plates for the TissueLyser tube adapter and homogenized on a TissueLyser II machine (QIAGEN, 85300) at 25 Hz, room temperature, for 5 min. After the first round of disruption/homogenization, we swapped the left and right adapters before initiating the second round. Swapping the adapters ensures that all samples receive uniform disruption and homogenization as samples closer to the TissueLyser are vibrated more slowly than those further away. After disruption/homogenization, the sample tubes were placed at room temperature for 3-5 min (this step helps dissolve lipid and membrane into the organic phase). Next, the lysed samples were transferred to 1.5 mL DNA loBind tubes (Eppendorf, 0030108051) that contained 200 µL chloroform, vortexed for 15 sec, and incubated at room temperature for 2-3 min. Samples were centrifuged at 12,000 x g, 4°C, for 15 min. For each tube, 350 µL total of aqueous phase (175 µL x 2) was transferred to another 1.5 mL DNA loBind tube that contained 350 µL 70% ethanol, followed by inverting the tubes 10 times to mix, and a brief centrifuge to collect all liquid. A total of 700 µL of each sample was transferred to an RNAeasy Mini spin column (reagent from QIAGEN, 74536), centrifuged at 10,000 x g, room temperature, for 30 sec (all subsequent wash steps use this centrifugation condition). The column was washed with 350 µL RW1 (reagent from QIAGEN, 74536) and incubated in 80 µL DNase I solution (prepared as instructed by the manufacturers) at room temperature for 15 min. To stop the DNaseI treatment, we added 350 µL RW1 directly to the column, which was then centrifuged and washed twice with 500 µL RPE buffer (reagent from QIAGEN, 74536) with a 2-min centrifugation step for the last RPE wash). RNA was eluted in 50 µL nuclease-free water (Invitrogen, 10977023) in a 1.5 mL DNA loBind tube, aliquoted, and stored at -80°C. RNA concentration was checked for all samples using a Thermo Fisher Varioskan LUX microplate reader μDrop plate (Thermo Fisher, N12391). Eight to ten RNA samples from each tissue were randomly selected to check RNA quality using an Agilent TapeStation 4200 (Agilent, G2991BA) and the TapeStation RNA ScreenTapes (Agilent, 5067-5576).

#### Liver

The tissues were first transferred from the collection tubes into 1.2 mL Collection Microtubes (QIAGEN, 19560) on dry ice in a 4°C cold room. A single autoclaved and pre-chilled (on dry ice) 5 mm stainless steel bead (QIAGEN, 69989) was added to each microtube. The microtubes were then quickly moved to wet ice, and 700 µL of QIAzol lysis reagent (QIAGEN, 79306) was added. Two rounds of homogenization were performed on a QIAGEN TissueLyserII at room temperature, 25 Hz, 5 min each. The lysate was transferred to new 1.5 mL DNA LoBind tubes, 200 µL chloroform (Fisher Scientific, C298-500) was added, and the tubes were vortexed for 15 sec and incubated at room temperature for 2-3 min. The subsequent RNA extraction protocol was performed as stated above. We note that good-quality RNA can be isolated using zirconium beads, which were used for the other tissues. This protocol was implemented due to a limited supply of reagents at the time. Lastly, the RNA from the liver samples of Cohort 1 was isolated separately from the other liver samples of Cohort 2.

#### Brain, gonads, and skin

All steps involving the RNAeasy Mini spin columns were performed on the QIACube HT robotic workstation (QIAGEN, 9001896) according to the manufacturer’s instructions, with the following program: 1) add 350 µL 70% ethanol to each sample aqueous phase in S-Block deep-well plate, mix, and transfer sample lysate into RNeasy 96 format vacuum columns (QIAGEN, 74104), 2) clear the columns using vacuum at 25 kPa for 3 min, 3) add 400 µL RWT buffer, 4) clear the columns using vacuum at 25 kPa for 1 min, 5) add 80 µL DNase I solution and incubate at room temperature for 15 min, 6) add 400 µL RWT, 7) clear the columns using vacuum at 35 kPa for 1 min, 7) add 400 µL 100% ethanol and incubate at room temperature for 2 min, 8) clear the columns using vacuum at 35 kPa for 1 min and then 25 kPa for 5 min, 9) add 45 µL nuclease-free water and incubate at room temperature for 4 min, 10) clear the columns using vacuum at 35 kPa for 1 min, 11) add 45 µL fresh nuclease-free water and 30 µL of the top elute fluid to the RNeasy 96-well plate and incubate at room temperature for 1 min, and 12) clear the columns using vacuum at 70 kPa for 2 min. The eluted RNA samples were aliquoted and stored at -80°C. We note that for some lipid-rich or debris-rich tissues, phase separation may be difficult (formation of the aqueous phase), making downstream processing challenging. To avoid this issue, for the ovary samples with high lipid content, QIAzol lysate was split into 2-3 aliquots after disruption/homogenization, topped off with QIAzol to 700 µL, and then processed individually until the column steps, before which they were pooled and passed over the same column. Several ovary samples were unfortunately not recoverable with this splitting method and were lost.

#### Bone

To facilitate tissue lysis, we ground the bone samples before the bead-beating step of the RNA extraction protocol. Briefly, an agate mortar, pestle, metal spatula, and a piece of aluminum foil were pre-chilled in liquid nitrogen. Bone samples were removed from -80°C (tubes stored in liquid nitrogen while awaiting processing) and placed on the chilled aluminum foil, which was then folded over in half to cover the bone sample. Covering the sample prevents larger chunks of the tissue from breaking apart and ‘flying’ out of the mortar. A pestle was used to press on the foil and grind the tissues into a powder. The powder was scooped using the pre-chilled spatula and placed into a 1.5 mL tube pre-chilled on dry ice. The bone ‘powder’ was stored at -80°C until RNA extraction.

#### Fat

Fat samples are prone to RNA degradation. We used the following modified RNA extraction protocol to preserve the RNA quality of fat samples. An agate mortar, pestle, and metal spatula were pre-chilled in liquid nitrogen. Fat samples were removed from -80°C (tubes stored in liquid nitrogen during await processing), transferred to the mortar, and ground to a fine powder with a rotating motion using the pestle. The powder was scooped using a pre-chilled spatula, placed into a 1.5 mL tube pre-chilled on dry ice, and stored at -80°C until RNA extraction. To extract the RNA from fat samples, we placed the frozen powdered fat samples on dry ice and added ∼100 µL of Zirconia/Silica beads to each tube. The samples were then transferred to wet ice, and 700 µL of QIAzol lysis reagent was added to each tube (the QIAzol-to-powdered tissue ratio was at least 2:1). The tissues were quickly homogenized on the TissueLyser II machine (the metal blocks from the tube holder adapter had been pre-chilled at -20°C) for 2.5 min at 30 Hz in a 4°C cold room. The tissues were incubated at room temperature for 5 min, centrifuged at 12,000 x g for 10 min at 4°C, and settled at room temperature for ∼2-3 min. The middle pink RNA layer was transferred into new tubes that contained 200 µL of chloroform, being careful not to aspirate the top lipid layer. Processing then proceeded in a similar manner to the other tissues.

#### Retina/RPE

The retina and retinal pigment epithelium (RPE) samples from each animal were dissected and processed together, as one tissue. RNA was isolated from the retina/RPE samples using the RNeasy Plus Micro Kit (QIAGEN, 74034) and following the manufacturer’s instructions. Briefly, 350 µL of Buffer RLT Plus was added to each sample, and the samples were homogenized by vortexing for 30 sec. The lysate was then applied to a gDNA Eliminator spin column and centrifuged at 8000 x g for 30 sec. The flow through was then combined with 350 µL of 70% ethanol, pipette mixed and then transferred to a RNeasy MinElute spin column. The column was centrifuged at 8,000 x g for 15 sec. The column was then washed with 700 µL of Buffer RW1 and then 500 µL of Buffer RPE, centrifuging at the previous settings after applying each wash and discarding flow-through. A final wash of 80% ethanol was applied to the column, and the sample tube was centrifuged for 2 min at 8,000 x g. Finally, the spin column membrane was dried by centrifuging the sample at full speed for 5 min. Then, the column was placed in a new 1.5 mL collection tube, 14 µL of RNAse-free water was applied to the membrane and centrifuged for 1 min at full speed to elute the RNA.

### Tissue RNA quality and sample dropout

We note that that two tissues have noticeable sample dropouts, including the retina/RPE and ovary (Extended Data Fig. 1b). This sample dropout could influence our downstream analyses (Spearman’s rank correlation and tissue aging clocks) given the lower sample size for these tissues. We note that the retina/RPE samples have different animal pooling strategies in the two cohorts and at different ages due to low RNA yield. These sampling and processing differences are reflected in the metadata contained in Supplemental File 2.

In the PCA plot (Fig. 1b), bone shows high sample variability compared to other tissues, possibly due to technical difficulties in preparing high-quality RNA from the bone. This high sample variability may influence our downstream analyses, leading to a low number of age-correlated genes and poor performance of the aging clocks.

### cDNA library generation and sequencing

cDNA libraries were prepared using a SmartSeq-based in-house protocol. Briefly, RNA samples were thawed on ice, and the concentration was measured using the Quant-iT RNA BR kit (Thermo Fisher, Q10213) on a Varioskan LUX Multimode microplate reader (Thermo Fisher, VL0000D0). RNA sample concentrations were normalized to 25 ng/µL, and 2 µL of each sample was used as input into the cDNA first-strand synthesis reaction. The resulting single-stranded library was amplified using 9 cycles. A portion of the full cDNA library volume (6 µL) was cleaned using Agencourt AMPure XP beads (Beckman Coulter, A63881) at a 0.7X ratio following the manufacturer’s guidelines, including two washes of 10.7 µL 80% ethanol (200 Proof, Gold Shield Distributors, 412804; diluted in nuclease-free water) and elution in 4.5 µL of nuclease-free water (Invitrogen, 10977023). The concentrations of the amplified cDNA libraries were measured using a Quant-iT dsDNA HS Kit (Thermo Fisher, 33120), and a subset of libraries were also measured on an Agilent TapeStation 4200 using a High Sensitivity D5000 ScreenTape (Agilent, 5067-5592).

Next, sequencing libraries were made using the Nextera XT DNA Library Preparation Kit (Illumina, FC-131-1096) and the IDT for Illumina DNA/RNA UD Index Sets A-D (Illumina, 2002713, 20027214, 20042666, 20042667), following the manufacturer’s instructions except for reducing all the reactions by half. Using half-volume reactions does not affect the performance of library preparation and conserves reagents for our large-scale experiment. The Illumina Index Sets A-D were converted into a 384-well format. Two library pools were ultimately generated, one of 322 samples and the other of 358 samples, with all the samples from the same tissue type assigned unique dual indices in the same library pool to reduce any batch effects. For tagmentation, 0.5 ng of the cDNA (2.5 µL total) was mixed with 5 µL TD buffer and then 2.5 µL ATM buffer from the Nextera kit, incubated at 55°C for 5 min and cooled to 10°C. To stop the tagmentation reaction, we added 2.5 µL of NT buffer and incubated the reaction mixture at room temperature for 5 min. The cDNA library was indexed and amplified for 12 cycles in a PCR reaction containing 10 µL of tagmented DNA, 5 µL of dual indices, and 7.5 µL NPM buffer. The amplified cDNA library (25 µL total) was split into two 12.5 µL aliquots, each purified using 22.5 µL of AMPure XP beads as described above. The aliquots were re-pooled after the first was eluted in 11 µL of Buffer EB (QIAGEN, 19086), such that the total elution volume was 10 µL. We performed most pipetting steps using the Dragonfly (SPT Labtech) or Mosquito HV (SPT Labtech) robotic liquid handlers to accelerate sample processing and maintain high pipetting accuracy. All steps requiring a thermocycler were performed on a 384-well plate thermocycler (BioRad). The concentration and quality of the library were measured using an Agilent TapeStation 4200 using a High Sensitivity D5000 ScreenTape (Agilent, 5067-5592). The experimental details for sequencing are provided in Supplemental File 2.

#### Shallow sequencing for normalization and quality assessment

To reduce sequencing depth variability across samples, we first performed shallow sequencing to more accurately determine the amount of each sample needed in a pooled library to achieve equal representation after sequencing. First, samples were pooled (1 µL per sample) across each row of each 384-well plate, resulting in 16 pools of 18-24 µL per plate. These 32 sub-libraries were quantified using a Qubit 1X dsDNA High Sensitivity Assay Kit (Thermo Fisher, Q33231) and analyzed on an Agilent 2100 Bioanalyzer (assays performed by the Stanford Protein and Nucleic Acid Facility) to determine the average library size. Then, two sequencing libraries (1 per 384-well plate) were generated by pooling the 16 sub-libraries per plate in an equimolar fashion, using the Qubit concentration and average library size. All samples from the same tissue type were kept in the same pool. The two pooled libraries were sequenced separately on an Illumina NextSeq 500/550 (Illumina) machine using two 150-cycle Mid Output v2.5 kits, 2 x 74 paired-end format (Illumina, 20024906). The on-instrument quality metrics, including Q30 and cluster densities, were in a suitable range for both sequencing runs.

We next ran the Bcl2fastq2 v2.20.0.422 program with 0.8 adapter trimming stringency on the sequencing run output files to generate FASTQ files for each pooled library. Each FASTQ file was processed using Trim-galore v0.4.5 to trim adapters and FASTQ v0.11.9 and multiqc v1.15 to assess sequencing quality. Total read counts were taken from the multiqc summary file ‘*mqc_fastqc_sequence_counts_plot_1.txt*’, looking only at the read 1 (R1) read counts (R1 and R2 read counts were comparable). We used the R1 read counts as input to calculate the volume of each sample needed for the deep sequencing libraries (2 pooled libraries as in the shallow sequencing), using a calculation template adapted from https://github.com/kalanir/CATechopooler/blob/master/COMET384_Seq7_Echo_Calculations.xlsx, and generated the pooled libraries based on the adjusted pooling numbers. There were 17 out of 697 samples omitted from the final deep sequencing libraries (680 samples remained) due to poor sequencing performance and library metrics.

#### Deep Sequencing

Each pooled library was sequenced by Novogene (Novogene, Beijing, China) on 2 (pooled library 1, which included 322 tissues) or 3 lanes (pooled library 2, which included 358 tissues) of an Illumina NovaSeq X 25B flow cell (2x150 bp paired-end) with 10% PhiX spike-in control for each lane, at a target sequencing depth of >40 million paired-end reads (20 million single-end) per sample. Novogene performed base calling, demultiplexing, and FASTQ file generation.

### Sequencing quality control and read mapping

Raw sequencing data (FASTQ files) were merged for each library pool (2 lanes for the pooled library 1 and 3 lanes for the pooled library 2) and checked for quality using Trim-galore v0.5.0. The processed reads were aligned to the African turquoise killifish reference genome downloaded from NCBI (Nfu_20140520, GCF_001465895.1) using STAR v2.7.10b^83^ with the default parameters. Out of all the sequenced samples, 14 samples had >90% of reads mapped to the genome; 252 samples, 80–90% reads mapped; and 178 samples, 75–80% mapped. Samtools v1.16.1^84^, with the parameters of MAPQ < 255 (‘samtools view -q255 -b’), was used to remove the reads mapped to multiple genomic regions. Next, we input the uniquely mapped reads into the ‘featureCounts’ program (with the default parameters) from subread v2.0.6^85^ to generate the read counts for each gene.

We detected three samples as outliers, which were removed from subsequent analyses: J6 (a liver sample), L21 (a testis sample), and H19 (a skin sample). Two samples (J6 and L21) were excluded because they had low total raw counts. One sample (H19) was excluded because it had low mapping performance. As a separate method, we used gene expression connectivity to detect outliers from the WGCNA package v1.73^86^. This method computes sample-to-sample correlations and derives network connectivity for each sample, then standardizes the connectivity scores, and finally identifies samples with Z-scores below -2 as outliers. Through this method, we verified these same three samples (J6, L21, and H19) as ‘outliers,’ validating their removal.

### Principal Component Analysis (PCA) and QC

All analyses of the Atlas RNA-sequencing data were performed in R v4.3.3 (apart from those described in the section ‘Calculation of Transcriptomic Age,’ which were performed in Python), and all the scripts are publicly available via GitHub (https://github.com/emkcosta/KillifishAtlas). First, to visualize the dataset quality, we created a *DESeqDataSet* object of all 677 samples using DESeq2 v1.42.1^87^. After filtering out genes for which the sum of the raw counts across all samples was 0 (15 genes), we applied the variance stabilizing transformation (‘vst’) on the raw counts stored in the whole-dataset-*DESeqDataSet* object and then visualized using the biplot function in the PCAtools package 2.14.0. The samples clustered nicely by tissue type along PC1 and PC2 (Fig. 1b).

The whole-dataset-*DESeqDataSet* object was then subset by tissue to generate individual tissue *DESeqDataSet* objects, which were stored in a list. To generate the PCA plot for a given tissue, we subset for the tissue and performed variance stabilization of the raw counts before running PCA as described above.

### Percent variance explained

We quantified the proportion of variance that could be explained by the covariates of sex, age, cohort, RNA extraction batch, RNA extractor, and the interaction of age:sex using the package variancePartition v1.33.11^88^ on a per tissue basis. First, ages were binned into six age groups (47-52 days, 75-78 days, 102-103 days, 133-134 days, 147-155 days, and 161-162 days), and age was modeled as the categorical variable ‘age_bin.’ Then, the TPM (transcripts per kilobase million) of each gene was generated for all samples. Next, the TPM count matrix was subset to include only the samples from a given tissue and prefiltered to only include genes with a TPM count > 0.5 in 80% of all samples of this tissue. For most tissues, the formula ∼ (1 | age_bin) + (1 | sex) + (1 | cohort) + (1 | RNA_batch) + (1 | RNA_extractor) + (1 | sex:age_bin) was used to explore the respective contributions of these variables to variance. For three tissues (bone, muscle, and fat), the formula ∼ (1 | age_bin) + (1|sex) + (1|cohort) + (1|RNA_batch) + (1 | sex:age_bin) was used, as the all of the RNA for these tissues had been extracted by one individual. For retina/RPE, the formula ∼ (1 | age_bin) + (1|sex) + (1|cohort) + (1 | sex:age_bin) was used, as all of the RNA for this tissue was extracted by the same individual in one batch.

The results of the variancePartition analyses for each tissue were saved in tabular format (as a CSV file) and plotted using the function plotVarPart.

### DESeq2 Differential expression analysis

To explore the age-sex interactions in our dataset, we performed differential expression (DE) analysis using DESeq2 on the tissue-specific *DESeqDataSet* objects (see ‘Principal Component Analysis (PCA) and QC). We first performed DE analysis using the design ‘∼ sex + age_bin + sex:age_bin,’ with ‘Female’ and the ‘age_bin1’ being the reference levels for sex and age_bin, respectively. The age_bin variable was modeled as a categorical variable (so as not to assume linearity), and we limited age_bin to bins 1-5 to focus on age bins for which we had sufficient sex balance (no female samples were collected in the 6th age_bin).

We next performed DE analysis between males and females in age_bin (1-5) using the design ‘∼sex’ (with ‘Female’ as the reference sex). For the sex-related differentially expressed genes (sex-DEGs) from this analysis, a positive log2-fold change occurs when the expression level for a gene is higher in males than females. A negative log2-fold change occurs when the expression level for a gene is higher in females than males. We plotted the prevalence of sex-DEGs (including both positive and negative DEGs) as a percentage of the total genes expressed in each tissue and each age_bin (Fig. 1e).

The analysis in Fig. 1e reveals that for each tissue for which variance partition analysis detects a contribution to variance by sex (either in the sex term or sex:age term), sex drives variance in a distinct manner.

### Identification of age-correlated genes

Age-correlated genes were identified on a per-tissue basis. First, a DESeq2 dds object was generated using the raw count matrix and sample metadata table subset for a given tissue and a given sex. Then, the raw count matrix was normalized using DESeq2’s ‘median of ratios’ method. To accelerate the identification of genes most correlated with age, we prefiltered this count matrix to only include genes that had a TPM count of > 0.5 in 80% of all samples in each tissue. These criteria exclude the genes with low counts, which are sensitive to noise in detection. After prefiltering, we used the processed normalized count matrix as input to calculate Spearman’s rank correlation between gene expression (normalized counts) and age, where age is the independent variable. A gene with an absolute value of Spearman’s rank correlation |*ρ*|>0.5 was considered an ‘age-correlated’ gene. While Spearman’s rank correlation captures monotonic behaviors, we employed other methods (see ‘Gene expression trajectory analysis’ below) to study the genes with other dynamics during aging (e.g., expressed in only one age, cyclic expression).

To identify age-correlated genes for both sexes combined, we used both male and female samples as one input for a given tissue before performing the same DEseq2 normalization, prefiltering (TPM count of > 0.5 in 80% of all samples for the given tissue), and Spearman’s rank correlation calculation.

To identify the age-correlated genes shared across tissues and both sexes, we first subset the atlas data by tissue, but we analyzed both male and female samples together when calculating Spearman’s rank correlation for each gene. Next, we found the intersection of the age-correlated genes (an absolute Spearman’s rank correlation of at least 0.5) in at least 6 tissues and plotted the Spearman’s rank correlation of each tissue as a heatmap (Fig. 2a). The Spearman’s rank correlations for each tissue are listed in Supplemental File 3 (sex-split) and Supplemental File 4 (sex-combined).

### Gene Set Enrichment Analysis (GSEA)

To perform GSEA^89^ on the age-correlated genes for each sex and tissue, we first calculated a ranked score for each gene by multiplying the Spearman’s rank correlation with the ‘-log10(p-value)’ and sorted all transcripts in descending order based on this score. Next, we used protein blast (best-hit protein with BLASTp E-value>1e-3) to identify the human ortholog for each killifish gene. The average of the ranked scores was calculated if multiple killifish paralogs were blasted to the same human gene. A killifish gene was removed if no human ortholog was found. Lastly, we ran the enrichment analysis via clusterProfiler v4.2.2^90,91^ and the Bioconductor annotation data package (org.Hs.eg.db v3.13.0). The p-values of the enriched pathways were corrected for multiple hypotheses testing using the Benjamini–Hochberg method (p.adjust). A Gene Ontology (GO) term (all three categories including biological process, cellular component, and molecular function, were tested) was considered significantly enriched if it had a value of p.adjust<0.05. The top GO terms significantly altered by age in both males and females were graphed as a dot plot in Fig. 1f and Extended Data Fig. 2. Extended Data Fig. 3 plots the GO terms significantly altered with age in only one sex and differed in the direction of change between the two sexes. The full GSEA data are listed in Supplemental File 5.

For selected GO terms (Fig. 1g), heatmaps were generated using Spearman’s rank correlations from males and females when the two sexes were analyzed separately (‘sex-split’). The above GSEA analysis outputs the human ortholog genes that drive each GO term. The gene lists of the same GO terms from males and females were merged, and the killifish genes corresponding to these human ortholog genes were identified (one human gene name can correspond to multiple killifish genes, and all the killifish genes were plotted). The heatmaps were generated using pheatmap v.1.0.12, with a defined scale from -1 to 1 (because the Spearman’s rank correlations do not exceed this boundary) and with the genes clustered.

### Hypergeometric Gene Ontology (GO) enrichment

We used the GOstats v2.68.0 packages for this analysis. The upregulated and downregulated genes shared across 5 tissues (derived from ‘sex-combined’ analysis and listed in Supplemental File 6) were separately used for the hypergeometric test implemented in GOstats v2.68.0. We used genes shared across 5 or more tissues to run this analysis because the gene set shared by 6 or more tissues was too small a set for this analysis. The background genes (‘universe’) were defined as all of the genes with a non-NA value for p.adjust for a given comparison. The full GO analysis results are given in Supplemental File 7.

### Gene expression trajectory analysis

Hierarchical clustering was performed on gene expression trajectories for genes expressed in all tissues. A gene was considered expressed if greater than 80% of the samples for a tissue type had a TPM of greater than 0.5. The intersection of expressed genes in each tissue resulted in 10,847 genes expressed in all tissues. For each tissue analyzed, the third age bin (102-103 days) was removed to avoid the lower sample number at this time point from driving the gene expression trajectory trend.

For each gene in each tissue, locally estimated scatterplot smoothing (LOESS) regression was performed to find a ‘trajectory’ for the Z-scaled normalized gene counts over age. These gene expression trajectories were then grouped into 10 clusters using hierarchical clustering (see the gene list for each expression cluster in Supplemental File 8). Genes that make up each cluster were then analyzed by Hypergeometric GO enrichment to identify enriched biological pathways. The full GO analysis results are given in Supplemental File 9.

### Identification of cell-type specific immune cell genes

The data exploration application (https://alanxu-usc.shinyapps.io/nothobranchius_furzeri_atlas/) associated with the publication^46^ was used to identify cell-type specific expression of immune genes. The ‘Bubbleplot/Heatmap’ tab was used to generate the gene expression dot plot for cell types (Extended Data Fig. 5a). The ‘CellInfo vs GeneExpr’ tab was used to generate UMAP plots with single gene expression overlayed (Extended Data Fig. 5f). The ‘Gene Coexpression’ tab was used to generate gene coexpression UMAP plots (Extended Data Fig. 5g). For all plots generated using this dataset for this publication, plots were downloaded as PNGs and edited slightly for figure clarity in Illustrator.

### *In situ* validation of the age-related gene expression changes

Tissues were collected from validation cohort animals (see ‘’African turquoise killifish husbandry’) and placed directly into ∼6 mL 4% paraformaldehyde (Santa Cruz Biotechnology, CAS 30525-89-4). Metadata for the animals used in each experiment is listed in Supplemental File 10. Samples were fixed for 16-24 h, washed with cold ∼12 mL nuclease-free PBS (Corning, 21-040-CM) for four 1-h washes, and then incubated in a nuclease-free methanol/PBS buffer series with each wash on ice for at least 10 min: 66% methanol (MeOH)/33% PBS, 100% MeOH, and 100% MeOH (Sigma-Aldrich, 3480-1L-R). Samples were then stored in fresh 100% methanol at -20°C until cryo-sectioning.

To prepare samples for cryo-sectioning, they were removed from -20°C storage and put through a reverse nuclease-free methanol/PBS buffer series to rehydrate the samples: 75% MeOH/25% PBS, 50% MeOH/50% PBS, 25% MeOH/75% PBS, and 100% PBS. Samples were incubated in 1 mL of each buffer for 15-30 min on ice. After the full methanol/PBS series, an additional wash in 1x PBS was performed for 15 min, and then samples were placed in 1 mL 30% nuclease-free sucrose solution (sucrose dissolved in nuclease-free 1x PBS, then filter-sterilized) and stored at 4°C overnight.

#### Tissue-specific embedding and sectioning strategies

The day after, samples were removed from the sucrose solution and dissected to prepare them for embedding. For each tissue, the dissection strategy was unique: kidney marrow “lobes” were dissected away from the muscle wall, and gut samples were cut lengthwise from the posterior to the anterior end to create a flat sheet. After dissection, samples were preincubated in the Neg-50 Frozen Section Medium (Fisher Scientific, 22-110-617) in individual wells of a 24-well plate (Corning, 353046) at room temperature for 10-15 mins. Before placing gut samples in Neg-50, they were gently rinsed in sucrose solution using a Pasteur pipette to wash away residual food debris from the lumen.

Following preincubation, each tissue type required unique embedding strategies in Neg-50. Kidney marrow lobes were embedded side-by-side, maintaining left-right and anterior-posterior orientation. Gut samples were rolled using the “Swiss roll” technique, with the anterior intestinal bulb’s luminal surface toward the center of the spiral and the posterior intestine toward the outside^92^. Samples were placed into a cryomold containing a thin (∼1-2 mm) sheet of frozen Neg-50 Medium on dry ice. Additional Neg-50 Medium was added, and the sample was left on dry ice to freeze fully. Frozen blocks were placed at -20°C until sectioning.

Samples were sectioned in batches by sex and age group so that the same section plane for each animal in the group was mounted on the same slide. All animals were given unique blinding IDs and deconvolved after quantification of mRNA spot count data. Samples were sectioned (30 µm) on a cryostat (Leica, CM3050 S), mounted on charged glass slides (Fisher Scientific, 22-037-246), and stored at -20°C until staining.

#### Hybridization chain reaction (HCR)

To validate mRNA expression from the atlas, a fluorescence *in situ* hybridization technique named hybridization chain reaction (HCR) was used^93^. The probes for each mRNA were designed using a custom-made Python script^93^, purchased from IDT (Newark, NJ, USA) as oPools and listed in Supplemental File 11. The following HCR amplifiers were purchased as solutions from Molecular Instruments (Los Angeles, CA, USA) and are listed in the format of ‘Amplifier-fluorophore’: B1-647, B3-546, and B5-488.

HCR was performed according to a protocol from Molecular Instruments (‘HCR RNA-FISH, fresh/fixed frozen tissue sections’). Briefly, tissue sections were equilibrated to room temperature from -20°C, rehydrated in 0.5-1 mL PBS for 5-10 min, and residual Neg-50 was gently washed off using PBS (‘Neg-50-free’). For the brain, the Neg-50-free sections were washed in 500 µL PBST (0.1% Tween-20 in nuclease-free PBS) four times, with 5 min incubation between each wash, and then incubated in 100-200 µL probe hybridization buffer (Molecular Instruments, buffer type: tissue section) at 37°C for at least 30 min (‘prehybridization’). To reduce the autofluorescence of the kidney samples, we incubated the Neg-50-free kidney slides in 1 mL 1x PBS and photobleached the slides under a strong LED light (‘RAYHOO 18W LED’, Amazon, B0CR1CHP7X) in a opaque chamber (cardboard box) at 4°C for at least 45 min. For the gut (and kidney optionally), the Neg-50-free sections were first baked at 60°C for 1 h in an *in situ* hybridization oven to increase adhesion between the tissue samples and the glass slides. After baking, the samples were rehydrated in 500 µL 100% ethanol, 500 µL 70% ethanol, and 500 µL 50% ethanol for 5 min incubation each. Next, the sections were post-fixed using 4% paraformaldehyde (diluted from 32% paraformaldehyde [Electron Microscopy Sciences 15714-S] in PBS) at room temperature for 15 min, followed by the four 500 µL PBST washes, with 5 min incubation between each wash, and then prehybridized.

After prehybridization, the buffer was removed, and 100 µL hybridization buffer (for each HCR probe, use 1 µL of the 0.5 pmol/µL stock per 100 µL hybridization buffer) was added to each slide, followed by 37°C incubation for 16-20 h. After hybridization, each slide was washed with 500 µL HCR probe wash buffer (Molecular Instruments, buffer type: tissue section), 500 µL 75% wash buffer (75% HCR probe wash buffer, 25% 5x SSCT), 500 µL 50% wash buffer (50% HCR probe wash buffer, 50% 5x SSCT), 500 µL 25% wash buffer (25% HCR probe wash buffer, 75% 5x SSCT), and 500 µL 5x SSCT (diluted from 20x SSCT [Ambion AM9770] with nuclease-free water) at 37°C with 15 min-incubation for each wash. Next, each slide was incubated in 200 µL HCR amplification buffer (Molecular Instruments, buffer type: tissue section) for 30 min – 4 h before switching to 100 µL amplification buffer supplemented with the fluorescent hairpin pairs (prepared according to the manufacturer’s instructions) for 20-24 h incubation at room temperature in the dark. Lastly, each sample was washed twice in 500 µL 5x SSCT/DAPI (10 µg/ml DAPI), with a 30 min incubation for each wash, followed by an optional 5-min 500 µL 5xSSCT wash. The slides were mounted with ProLong Gold Antifade reagent (Thermo Fisher, P36934) and sealed with nail polish.

The slides were imaged on a Zeiss LSM900 confocal laser scanning microscope (Zeiss) equipped with Zen 3.0 (blue edition) software, Zeiss Plan-Apochromat 40x/1.4 oil objective, and Zeiss Immersol oil 518F (Zeiss, 4Y00-R0DY-1007-3VF3) as an immersion medium. The imaging conditions were the following: 9-slice z-stacks with a step size of 0.75 µm; the Alexa Fluor 546 channel (laser at 1%, detector gain: 775V, detector offset: 256, detector digital gain: 1.0); the Alexa Fluor 488 channel: (laser at 2.5%, detector gain: 650V, detector offset: 256, detector digital gain: 1.0); the Alexa Fluor 647 channel (laser at 8.0%, detector gain: 650V, detector offset: 512, detector digital gain: 1.0); and DAPI (laser at 0.5%, detector gain: 650V, detector offset: 256, detector digital gain: 1.0). Four fields of view per tissue section and four animals per condition were imaged. All images were taken in comparable regions across biological replicates, specifically along the caudal-rostral axis of the ‘Swiss roll’ for the gut (using individual villi as landmarks), and along the caudal-rostral axis for the kidney (interstitial and kidney tubule epithelial regions).

#### Quantification of HCR Images

All samples were blinded and randomized after tissue harvest. Each sample was assigned a sample ID, which was used for sample processing and imaging, and the sample information was not revealed until after image quantification. To quantify *IGF2BP3* (LOC107383282), *LOC107373777* (*ncRNA-3777*), and *irf4a* (*LOC107383908*) mRNA levels, we first performed maximum-intensity projection in the z-direction for all images using a FIJI^94^ macro script (z-planes 4-6 were used for *IGF2BP3* and *ncRNA-3777* and all 9 z-planes were used for *irf4a*). Max-projected images were then loaded into QuPath software (v.0.5.1, https://qupath.github.io/) to quantify mRNA spots. First, the cells were segmented using a nuclear mask created based on the DAPI signal (DAPI threshold: 3000; sigmaMicrons: 1.5; minAreaMicrons: 10.0; maxAreaMicrons: 400.0), and then an expansion of 10 µm from the DAPI mask was used as the cell boundary. Detection of red blood cells, which have strong autofluorescence in all channels, and cells located in the kidney tubule regions were manually removed to avoid false positive subcellular spot detection. Next, the QuPath subcellular detection function was used to detect each type of mRNAs using specific parameters. Because the gut images have highly variable background signals, each image requires a separate threshold to detect signal from noise for counting the *IGF2BP3* and *ncRNA-3777* mRNA spots. To consistently distinguish signal from background, for each fluorescent channel, we plotted the distribution of the maximum signal of each cell (k), found the mean value of k, and defined ‘signal’ to be at least 0.5 standard deviations above the mean of k in the image. This method matches with manual counting well. The kidney images have mostly consistent backgrounds, so the same QuPath detection parameters were applied to most of the kidney images, but a subset of the kidney images needed different parameters to accommodate a high background (see Supplemental File 12 for full parameter record and results). For *irf4* quantification, only cells located in the interstitial region are counted. After QuPath detection, every cell was visually inspected to check the detected spots matched with manual counting. A small number of false positive spots were manually removed (these spots usually occur in regions overlapping with red blood cells). The number of a specific type of mRNA per cell and the cell number of each image were recorded (Supplemental File 12).

We reported the average mRNA counts per cell for each animal. For this calculation, we first found the total number of mRNA spots by summing all the cells across four fields of view imaged for an animal. The average number of mRNA counts per cell was calculated by dividing the total number of mRNA counts by the total number of cells summed across the four fields of view. To compare the young and old differences in the average mRNA counts within one sex, statistical significance was calculated using the Mann-Whitney test. Two-way ANOVA (sex, age, sex-age interaction) was used when analyzing both sexes together.

### Cell dissociation and flow cytometry of killifish head kidney

Animals were randomly selected from the validation cohorts to use for flow cytometry experiments. Three batches of young and old animals were processed for head kidney flow cytometry for males and two batches were used for females. Experimental metadata are documented in Supplemental File 13.

Fish were anesthetized for 1.5 min in an ice slurry made using system water. Once operculum movement slowed, and the fish was no longer responsive to touch, the animal was dissected and transcardially perfused with 10 mL of ice-cold 0.25 M EDTA solution (Fisher Scientific, AAJ15694AP) in 1x PBS (Thermo Fisher, 10010049) as described above. Following perfusion, head kidney tissue was carefully dissected from the body wall and placed in 5 mL of ice-cold fetal bovine serum (FBS) (Fisher, 50-152-7067) in a well of a 12-well culture plate (Cell Treat, 229111). This process was repeated until all animals in the batch were perfused and dissected.

Single-cell suspensions from head kidney tissue were prepared for flow cytometry using a non-enzymatic dissociation protocol adapted from zebrafish^95^. Kidney marrow in FBS was pipetted 50 times with a 5 mL serological pipette to mechanically dissociate the tissue. The digestion mixture was then applied to a 100 µm Cell Strainer (Fisher Scientific, 07-201-432) sitting atop a 50 mL conical tube (Fisher Scientific, 1443222). Tissue clumps remaining on the mesh were gently triturated using the plunger of a 1 mL syringe (Fisher Scientific, 14-826-88), and then 5 mL of SM Buffer (5% FBS in 1x PBS) was used to wash the well of the 12-well plate and the 100 µm strainer mesh. Filtered cells were then pelleted (400 x g, 4 min, 4°C), and the supernatant was removed using a 10 mL serological pipette until about 200 µL remained. The pellet was then resuspended in 5 mL of SM buffer by pipette 5 times and then was applied to a 40 µm Cell Strainer (Sigma-Aldrich, CLS431750-50EA) on top of a 50 mL tube. The strainer was then washed with 2 mL of SM buffer, and cells were pelleted once more using the previous conditions. The supernatant was again removed (leaving about 100 µL of SM Buffer), and pellets were resuspended using 500 µL of additional SM Buffer. The cell suspension was moved to a 1.5 mL Low-Adhesion Tube (USA Scientific, 1415-2600) and centrifuged at 400 x g for 2 min at 4°C. Finally, the supernatant was removed until 200 µL remained. Cells were resuspended, and about 5-10 min before the sample loading onto the cytometer, the live/dead stain 7-AAD (BD Biosciences, 559925) was added. Right before loading on the cytometer, the cell suspension flowed through the 35 µm strainer mesh cap of a 5 mL round-bottom FACS tube (Corning, 352235). Then, the sample was loaded for analysis and/or sorting on a Sony MA900 Cell Sorter (nozzle size: 100 µm, flow rate: 4).

Gates were drawn to exclude debris and to capture live, single cells. Then, gross populations of immune cells (erythroid, myeloid, lymphoid, progenitor, all leukocytes) were identified by side-scatter and forward-scatter based on a protocol developed for zebrafish^51^. The myeloid: lymphoid ratio was calculated by dividing the total number of myeloid cells by the total number of lymphoid cells for each sample. For all cytometry plots used for quantification, a total of 50,000 events were recorded. Cytometric analysis was performed using FlowJo version 10.10.0.

### Bulk RNA-sequencing of FACS-sorted cells

Five males of various ages (67, 88, and 201 days) were used to test the gating strategies for sorting different populations of kidney-dissociated cells by FACS. The number of cells collected per sample is listed in Supplemental File 14. Cells were sorted into 350 µL Buffer RLT Plus (reagent from the QIAGEN RNeasy Plus Micro Kit [QIAGEN, 74034]) containing β-mercaptoethanol (10 µL β-ME per 1 mL Buffer RLT Plus), briefly vortexed for 30 sec, and then frozen immediately on dry ice. For sorted volumes exceeding 500 µL, additional Buffer RLT Plus was added to the sorted cells before vortexing and freezing at a ratio of 350 µL Buffer RLT Plus for each additional 100 µL of cytometer sheath fluid. Frozen homogenates were stored at -80°C until RNA extraction.

Bulk total RNA extraction of the sorted cell populations from flow cytometry was performed using the QIAGEN RNeasy Plus Micro Kit (QIAGEN, 74034) according to the manufacturer’s instructions. The frozen homogenates of sorted cells were thawed on ice for 30 min. Once completely thawed, homogenates were briefly centrifuged, then applied to a gDNA Eliminator spin column and centrifuged at 10,000 x g for 30 sec. The flow-through was then added to a DNA LoBind tube (Sigma-Aldrich, 022431021) containing an equal volume of 70% ethanol. This process was repeated until all the remaining volume of homogenate was passed through a gDNA Eliminator spin column and mixed with an equal volume of 70% ethanol. Then, samples were pipette mixed, transferred to a RNeasy MinElute Spin Column, and centrifuged at 10,000 x g for 15 sec, discarding the flow-through. This step was repeated until the entire ethanol-homogenate mixture was applied to the RNeasy MinElute Spin Column. The column was then washed twice, first with 700 µL of Buffer RW1 and then 500 µL of Buffer RPE, centrifuging at 10,000 x g for 15 sec and discarding the flow-through each time. A final, longer wash was performed with 80% ethanol, after which the spin column was centrifuged for 2 min at 10,000 x g. Then, the spin column was transferred to a new collection tube and dried by centrifuging at 12,000 x g for 5 min. Finally, the column was transferred to a new 1.5 mL collection tube, 14 µL of RNAse-free water was applied to the membrane and then was centrifuged for 1 minute at 12,000 x g to elute the RNA. RNA was quantified using the Quant-iT RNA BR kit (Thermo Fisher, Q10213) on a Varioskan LUX multimode microplate reader, aliquoted, and stored at - 80°C.

cDNA and library synthesis were performed using a modified in-house SmartSeq2 pipeline similar to as described above for whole tissues (see section titled ‘cDNA library generation and sequencing’), with a few modifications to accommodate lower input concentrations of RNA. First, the single-stranded library was amplified using 16 cycles. Next, tagmentation was performed using 0.1 µL of the Illumina Tn5 enzyme (Illumina, 20034198), 0.26 µL of nuclease-free water, and 0.64 µL of 2.5X TAPS-PEG crowding agent per sample. The crowding agent was prepared by combining filtered 40% w/w PEG 8000 (Promega, V3011) 1:1 v/v with 5X TAPS-MgCl_2_ (3 mL of 0.5M TAPS-NaOH pH 8.5 [Boston Bioproducts, BB2375] combined with 750 µL of 1M MgCl_2_ [Sigma-Aldrich, M1028] and 26.25 mL of nuclease-free water, and adjusted to pH 8.4). The resulting tagmented library was amplified for 12 cycles using the Kapa enzyme (KAPA Hifi PCR kit, Kapa Biosystems, KK2102).

RNA-seq analysis (quality control, mapping, count generation, DESeq2 analysis) and plotting were also performed as in the atlas dataset, except using a different expression cutoff from the atlas: in this case, genes with at least one count in at least one sample were retained (rather than requiring at least 80% samples having at least one count). The Principal Component Analysis plot was generated using the ‘plotPCA’ function in the DESeq2 (v.1.34.0), and the heatmap by ‘pheatmap’ (v.1.0.12) with the parameters: rows and columns clustered, Z-scale for each gene, and capping z-scale at -2 and 2 (any value below -2 or above 2 were assigned as the lowest color or the highest color, respectively).

### Calculation of Transcriptomic Age

#### Training Tissue Clock Models using BayesAge 2.0

All computation relating to the tissue clocks were performed using Python (v3.11.11) in a series of Jupyter notebooks run in the free version of Google Colaboratory. Transcriptomic age was calculated using the published method BayesAge 2.0^71^. This method utilizes a Bayesian framework to estimate the most likely transcriptomic age of a sample (‘tAge’) and employs locally weighted scatterplot smoothing (LOWESS) regression to model the nonlinear dynamics of gene expression, enabling age prediction between 47 to 163 days of age at day-level resolution.

Before training tissue-specific models, we first preprocessed the raw gene expression matrix. Raw gene expression counts were normalized using frequency count normalization, whereby raw counts were transformed into relative frequencies by dividing the raw count for each gene by the total read count for the sample. Next, LOWESS regression was used to fit a trend for each gene across age.

After preprocessing, we performed model training. We employed Leave-One-Sample-Out Cross-Validation (LOSO-CV) to separate our dataset into training and test sets: For each tissue clock, we separated the tissue dataset of size N into a training set of N-1 samples and a test set of one sample (‘left out’). For each training-testing group, we first trained a reference matrix by taking the gene frequency counts for each gene for the N-1 samples in the training set, computed a LOWESS regression fit, and performed feature selection for enhanced biological interpretation. To select features of interest, we calculated the Spearman’s rank correlation between gene frequency and age for each gene. A set of genes (groups of 5, 10, 15, etc., up to 50 genes were iteratively tested) with the highest absolute Spearman’s rho were used for age prediction. It is important to note that each time LOSO-CV is performed, the identities of the top Spearman’s rank correlated genes may differ, as leaving a different sample out may slightly alter the relationship between age and gene expression. The resulting trained matrix stores the predicted gene frequency levels using LOWESS fit across age and the Spearman’s rho values for each gene in the dataset for these N-1 samples.

Next, we performed age prediction for the ‘left out’ test sample. We selected a given number (M) of top Spearman’s rank correlated genes (different M values were included during testing), and for each gene, we computed the probability of observing the gene expression for that gene for a particular age, assuming a Poisson distribution. The probability for the age-related gene state is given by:

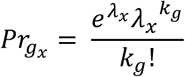

where:

*x*: specific age

*λ_x_*: expected gene expression count at age *x*

*k_g_*: observed gene expression count for the test sample, *ϕ*

The expected gene expression count was derived from the frequency-normalized trained reference matrix, and the observed gene expression count came from the observed raw counts of a particular gene from the M genes for the test sample.

Then, for each age *x*, the probability of the test sample is a given age was the product of the individual probabilities for each gene.

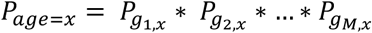

The age prediction (tAge) for the test sample was then found by computing an age-likelihood distribution and finding the maximum likelihood age.

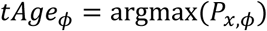

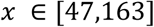

To avoid numerical underflow errors during computation, we replaced the product of individual gene probabilities at a given age with the sum of logarithms of these individual gene probabilities and found the maximum likelihood age from this distribution. This preserves the numerical relationship and avoids Python rounding errors.

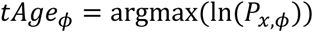

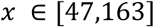

We repeated this process, leaving out a different sample from the tissue dataset until each sample had been tested. After this process, we obtained the age predictions for each of the samples in our tissue dataset. Performing LOSO-CV with different gene set sizes (M) informed us of the optimal M that corresponds to the most concordance between chronological and predicted age, and we called this optimal condition for a tissue clock the ‘optimal clock’ using the BayesAge model. We calculated the Pearson correlation (r), Coefficient of Determination (R^2^), and Mean Absolute Error (MAE) using the Python scipy (version 1.13.1) package to evaluate model performance. The results for LOSO-CV for BayesAge are summarized in Supplemental File 15.

#### Comparison of BayesAge 2.0 to Other Models

The primary advantages of BayesAge 2.0 over other common modeling strategies for ‘omics’ data, such as Elastic Net regression (EN) and Principal component regression (PC-R), are that it 1) reduces data overfitting, 2) does not require extensive hyperparameter tuning (a time-intensive process), and 3) has enhanced biological interpretability due to feature pre-selection. We developed EN and PC-R models for each Atlas tissue dataset to benchmark BayesAge 2.0 model performance.

To perform age prediction using Elastic Net regression, we used DESeq2 normalized counts. We z-scaled the gene expression data using the StandardScaler function in the scikit-learn Python module (version 1.5.2). Elastic Net is a linear regression model that combines Lasso (L1) and Ridge (L2) regularization. To optimize model performance, Elastic Net requires tuning of two hyperparameters, *α* and *λ*, which control the trade-off between L1 and L2 regularization and the strength of the regularization, respectively. To implement hyperparameter tuning, we performed a parameter grid search using the GridSearchCV function from scikit-learn for the parameter *λ* (called alpha in scikit-learn’s implementation of Elastic Net) and *α* (called l1 ratio in scikit-learn). This search was performed in two steps: first, we tested alpha values from 1e-5, 1e-4, 1e-3, continuing up to 100, and l1 ratio values from 0 to 1 in step sizes of 0.1. The maximum number of iterations was set to 10,000 for most tissue clocks (except for the brain, for which it was set to 30,000). We implemented LOSO-CV for each combination of alpha and l1 ratio parameters using the LeaveOneOut function in scikit-learn. To evaluate model performance, we used MAE. Once optimal parameters were identified, the second step involved increasing the maximum iteration number of 100,000 for all tissues to ensure objective function convergence and to finalize age predictions. The ‘optimal’ tissue clock using Elastic Net uses the optimal parameters for *α* and *λ* derived from hyperparameter tuning. For these analyses, random seeding was set to 42 to ensure reproducibility. The results of hyperparameter tuning and LOSO-CV are summarized in Supplemental File 16.

As in our implementation of Elastic Net, we used the DESeq2 normalized counts to implement Principal Component regression and then scaled the data using the StandardScaler function. PC regression is a regression technique that combines principal component (PC) analysis and linear regression, fitting a linear regression model using a subset of the PC’s as predictors. To perform PCA analysis, we used the PCA function in scikit-learn. We implemented LOSO-CV for each PC number from 5 to 20 in steps of 5 using the cross_val_predict function in scikit-learn and we evaluated model performance using MAE. The ‘optimal’ tissue clock using PC regression occurs when a PC number is identified that maximizes the coefficient of determination and minimizes MAE. For these analyses, random seeding was set to 1 to ensure reproducibility. Results of LOSO-CV are summarized in Supplemental File 17.

To evaluate the performance of BayesAge 2.0 in comparison to other models, we compared the residuals from BayesAge 2.0, to EN and PC-R by computing the residuals in two ways, first as the difference between the predicted age and the line of best fit and second as the difference between predicted age and true chronological age for each sample. We found that BayesAge 2.0 has the lowest bias in residual distribution (Extended Data Fig. 6), suggesting the validity of using BayesAge 2.0 for our modeling.

#### Age Prediction in Other Datasets

To demonstrate the generalizability of our tissue-specific clocks to other datasets, we performed age prediction in an additional published RNA-sequencing dataset^22^, which is a liver transcriptomic dataset (‘AL/DR’) from male and female killifish fed on *ad libitum* (AL) and dietary-restriction (DR) diets from sexual maturity (4 weeks) to 9 weeks of age.

We performed age prediction in this query dataset using the three different machine learning models described above: BayesAge 2.0, Elastic Net regression, and Principal Component regression. For male killifish liver samples from the query dataset, we used the trained reference matrix from only male liver samples in the Atlas dataset (male-specific liver clock) for age prediction. For female killifish liver samples in the AL/DR dataset, we used the female-specific liver clock to make predictions.

First, to perform age prediction in a query dataset using BayesAge 2.0, we trained a reference matrix containing all N samples in the Atlas dataset for the tissue type of the query samples. Then, we computed the predicted age (tAge) for the sample in the query dataset using an age-likelihood distribution and finding the maximum likelihood age. For each gene-wise probability, the trained reference matrix serves as the source of expected gene counts, and the raw expression matrix of the query dataset serves as the source of observed gene expression. The gene number M used in the models varied from 5 to 100 (at an increment of 5 genes), and the results for each M were reported.

To perform Elastic Net regression age prediction in a query dataset, we used the optimal parameters for *λ* and *α* as derived from hyperparameter tuning. Hyperparameter tuning was performed as described in ‘Comparison of BayesAge 2.0 to Other Models’, separately for the male and female liver clocks. These optimal parameters were then used to retrain the male- or female-specific models using the atlas data, and the query dataset was used as testing data to perform age prediction using the optimized model. The non-zero coefficients, which are the genes that the model uses to perform age prediction, are reported, as well as their coefficient values (weights).

For age prediction using Principal Component regression (PC-R), PC-R models were made as described in ‘Comparison of BayesAge 2.0 to Other Models’. LOSO-CV using atlas samples was performed separately for male and female liver clocks. After model training, the query dataset was used as the testing data and the optimal PC number for age prediction in the test set was identified as where the Mann-Whitney U (MW) test p-value stabilized.

The predicted age data for all three models are listed in Supplemental File 18.

As a measure of effect size between predicted ages of control and treated animals, we computed ΔtAge in two ways: as the difference in medians or means between control and treatment groups. We assessed whether the predicted ages of the control (young, AL, or wildtype) and treatment groups (old, DR, or mutant) differed using a few measures: a comparison of distribution shape Kolmogorov-Smirnov (KS) test, a comparison of distribution central tendency Mann-Whitney U test, and finally a simple calculation of percent overlap of the age prediction distributions.

---

**Extended Data Figure 1:**
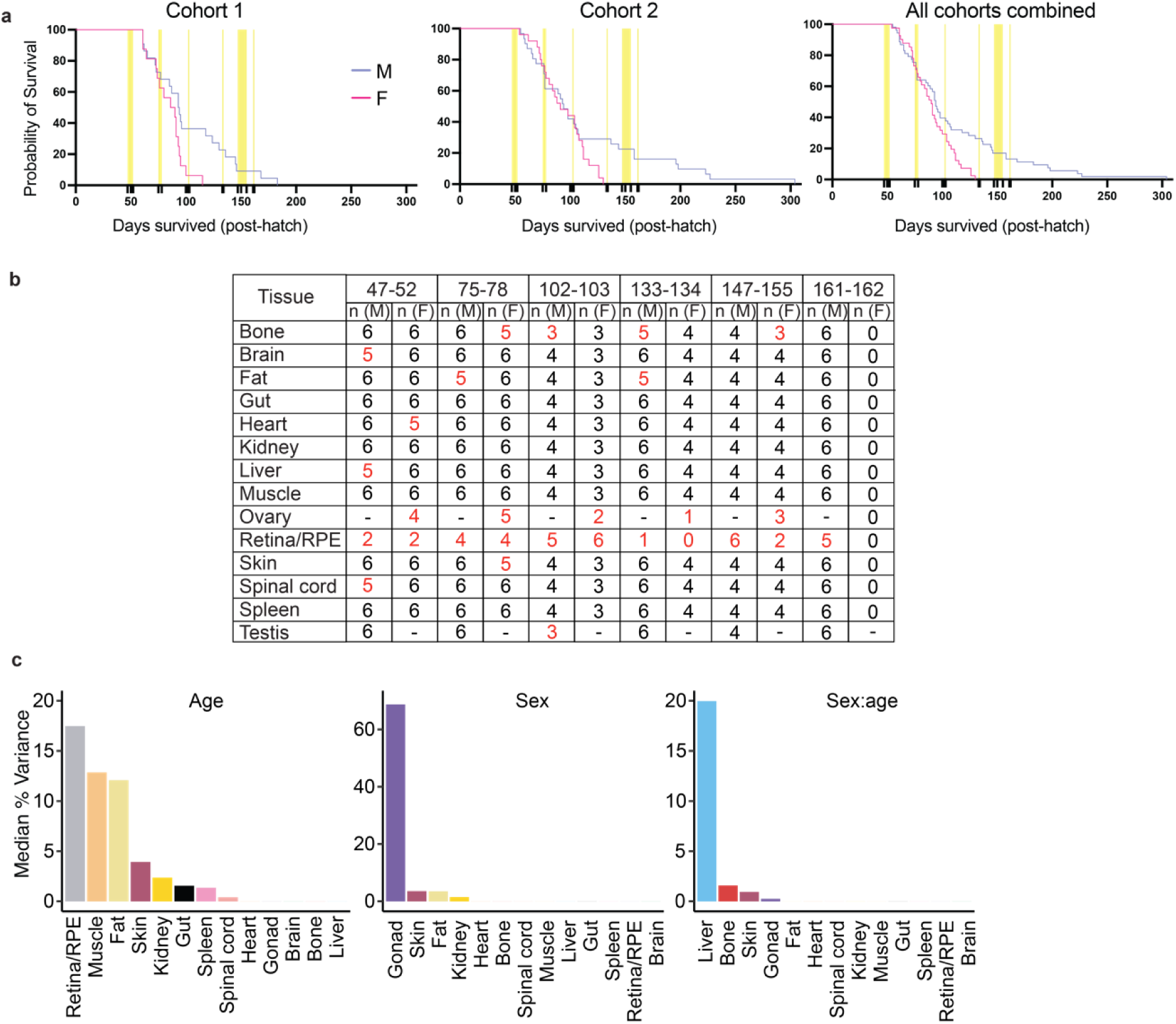
Metadata for the multi-tissue killifish transcriptomic aging atlas. (a) Kaplan-Meier survival curves for the two cohorts (left, middle) from which samples for RNA-sequencing were derived (left, 19 females, 24 males; middle, 31 females, 33 males). On the right is the survival curve for both cohorts combined (50 females, 57 males). Blue, male survival curve; red, female survival curve. Yellow and additional ticks on x-axis, sample collection windows. F, female; M, male. (b) Number of samples analyzed for each tissue, sex, and age group in this study. The red numbers denote incidences of sample dropout. ‘—’ indicates ‘not applicable.’ (C) Bar plots of the median percent variance explained across all genes expressed in an tissue for the covariates of age (left), sex (middle), and the interaction term sex:age (right).

**Extended Data Figure 2:**
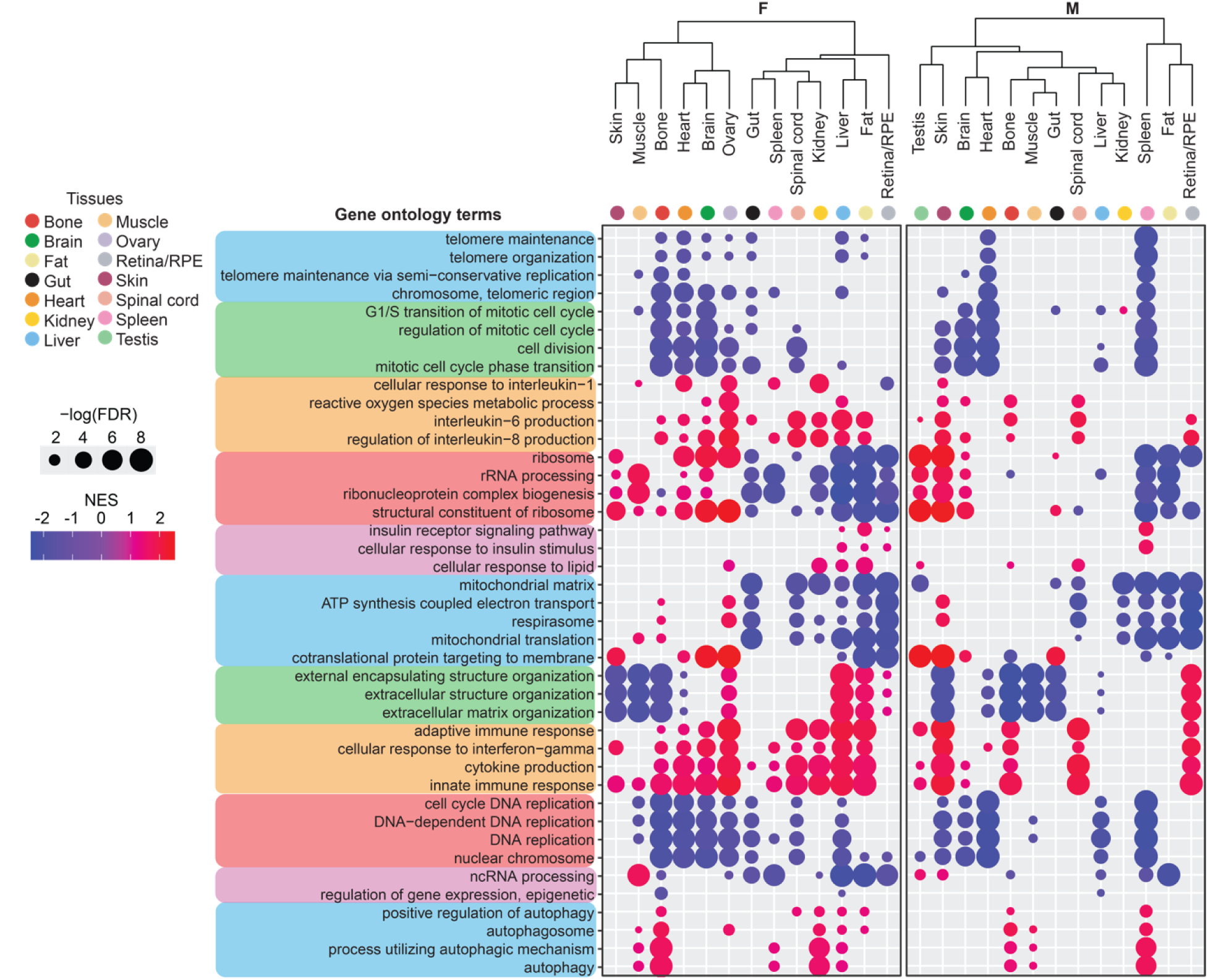
Cross-tissue pathways enriched for the genes correlated with age. Male (M) vs. female (F) gene set enrichment analysis (GSEA) results, identifying the shared or unique pathways enriched for the genes upregulated or downregulated with age in the 13 tissues. For females and males, separately, tissues are clustered by similarity of enrichment as calculated by the product of the NES and –log(FDR). NES, normalized enrichment score. Dot size, –log10 of the adjusted p-value (i.e., false discovery rate [FDR] after multiple hypotheses testing).

**Extended Data Figure 3:**
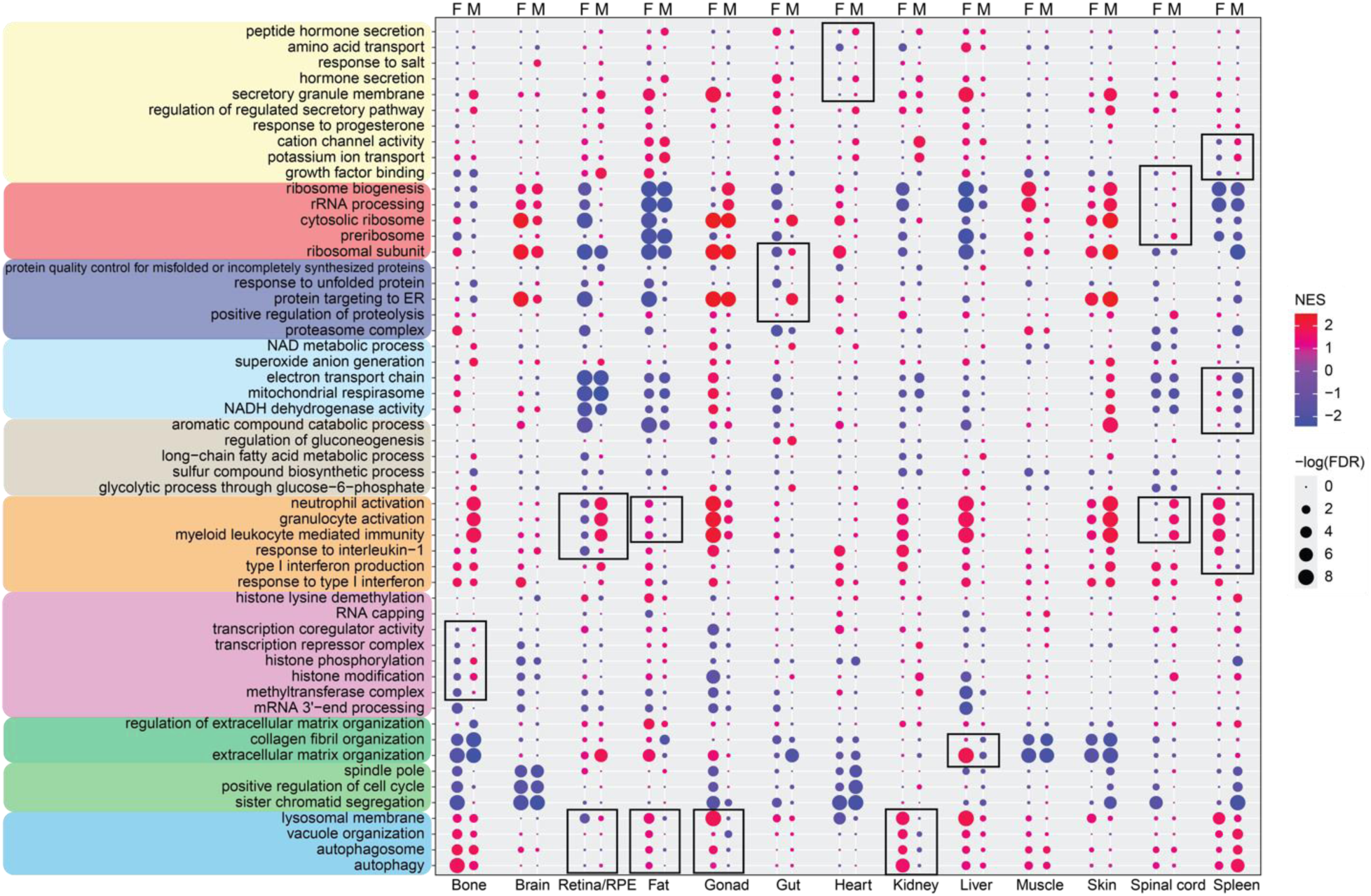
Sex-specific pathways enriched for the genes correlated with age. Male (M) vs. female (F) gene set enrichment analysis (GSEA) results in the 13 tissues, identifying the GO terms showing opposite signs of upregulation or downregulation with age in the two sexes, and those for which the change with age is significantly in only one sex (‘sex-divergent’). NES, normalized enrichment score. Dot size, –log10 of the adjusted p-value (i.e., false discovery rate [FDR] after multiple hypotheses testing). Boxes indicate the main sex-divergent GO terms in each tissue.

**Extended Data Figure 4:**
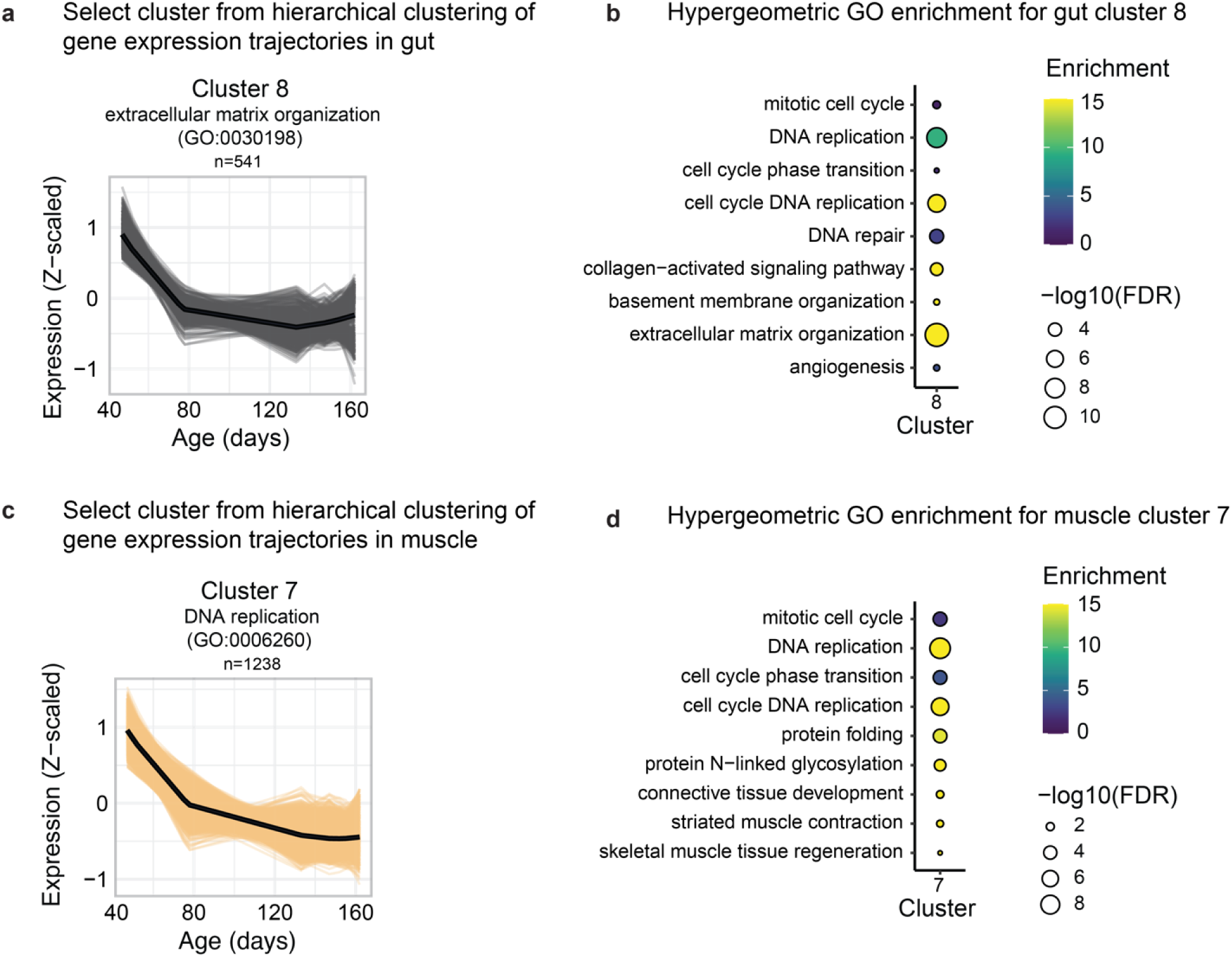
Tissue-specific gene expression dynamics for the gut and muscle. (a) Hierarchical clustering of the gene expression trajectories for the gut (sex-combined), highlighting cluster 8. The average trajectory for the cluster is depicted by the black line. The most highly significant GO term from Hypergeometric GO enrichment (terms related to Biological Processes) is listed, as well as the number of genes making up the cluster. (b) Hypergeometric GO enrichment (terms related to Biological Processes) for the genes in gut cluster 8. Select significantly enriched (adjusted p-value < 0.05) GO terms for each cluster are plotted. Dot color represents the enrichment score of each GO term, with the maximum value of the scale adjusted to 15 to improve color resolution of GO terms with lower enrichment. Dot size, –log10 of the adjusted p-value (i.e., false discovery rate [FDR] after multiple hypotheses testing). (c) Hierarchical clustering of the gene expression trajectories for the muscle (sex-combined), highlighting cluster 7. The average trajectory for the cluster is depicted by the black line. As in panel a, the most highly significant GO term from hypergeometric GO enrichment is listed, as well as the number of genes making up the cluster. (d) Hypergeometric GO enrichment for muscle cluster 7, analysis conducted as in panel b.

**Extended Data Figure 5:**
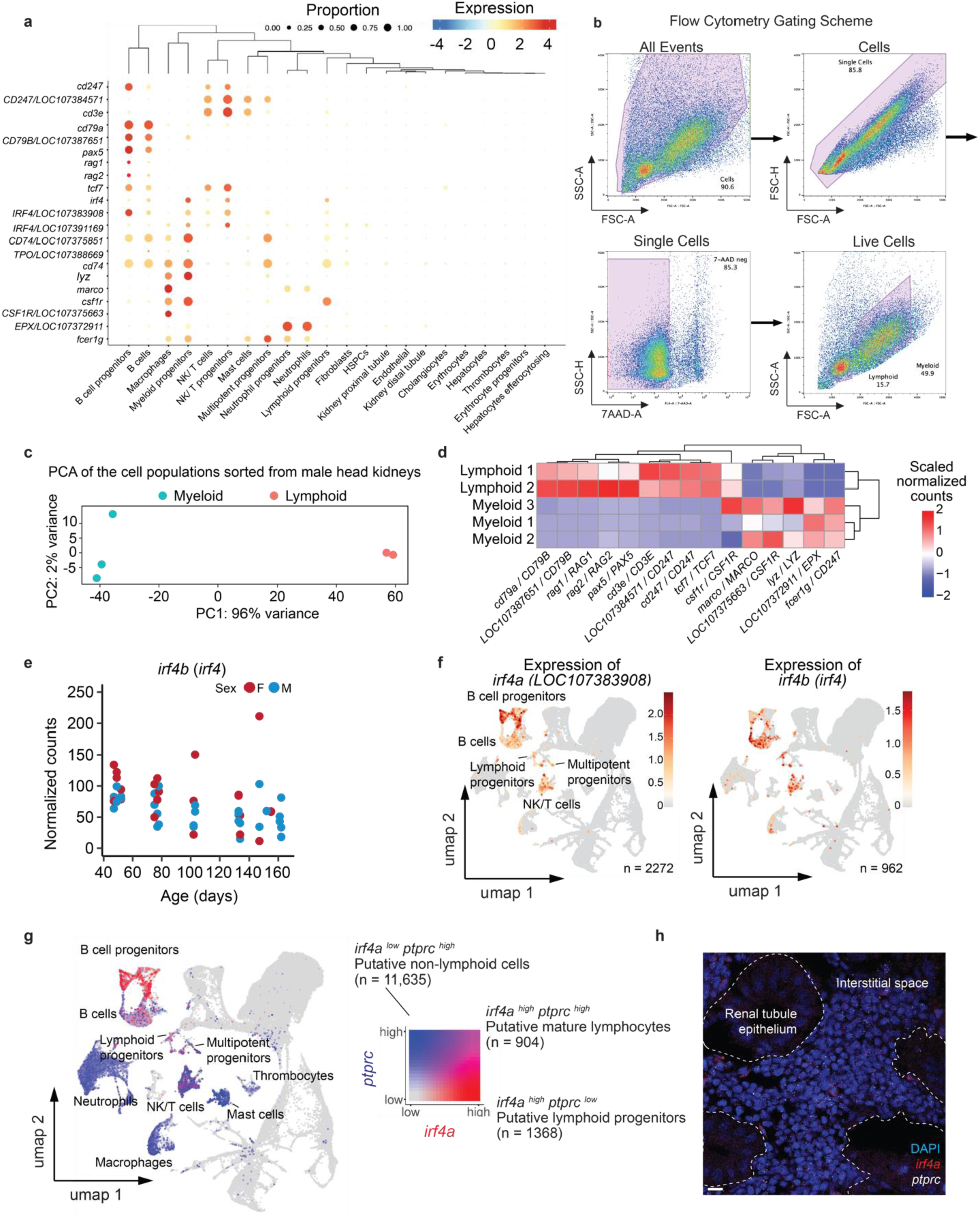
The aging killifish kidney marrow changes in gene expression and cell-type composition. (a) Dot plot of gene expression for genes in Fig. 4b, showing cell-type specific enrichment. Dot color indicates the level of expression, and dot size indicates the percentage of cells expressing the gene. (b) Flow cytometry gating scheme, showing representative gating workflow from raw event data to live cells. (c) Principal Component (PC) Analysis of the dissociated male head kidney cell populations that were FACS-sorted based on the gating strategy as in panel b. Each dot is an individual animal (myeloid population: 3 fish; lymphoid population: 2 fish). These males were harvested from different ages (67, 88, and 201 days) to test whether the gating strategy can be applied to different age groups. (d) Heatmap showing the expression of myeloid and lymphoid cell type-specific markers (see panel a), clustered by samples. The expression of each gene is plotted as Z-scaled, DESeq2-normalized counts. (e) Scatterplot of the counts normalized by DESeq2 for *irf4b* (killifish gene name: *irf4*) in the head kidney transcriptome of the atlas dataset. Each dot is the expression of *irf4b* in an individual sample. Red, female (F). Blue, male (M). (f) UMAP (uniform manifold approximation and projection) plots of data from a killifish single-cell RNA-sequencing tissue atlas^46^, with overlayed expression levels for *irf4a* (left) and *irf4b* (right). (g) Co-expression UMAP showing the expression level of *irf4a* and *ptprc*. Data were derived from the tissue atlas^46^. *The irf4a^high^ ptprc^low^* cells are red (1368 cells in the source dataset), *irf4a^low^ ptprc^high^* cells are blue (11,635 cells), and *irf4a^high^ ptprc^high^* cells are purple (904 cells). (h) Example single-z-plane HCR image of young male head kidney tissue, with cross section of renal tubule epithelium encircled by white dashed lines. Outside of these white dashed boundaries is the interstitial space, where hematopoietic tissue resides. Quantification of the *irf4a* transcripts was performed for the interstitial space. Scale bar, 10 µm.

**Extended Data Figure 6:**
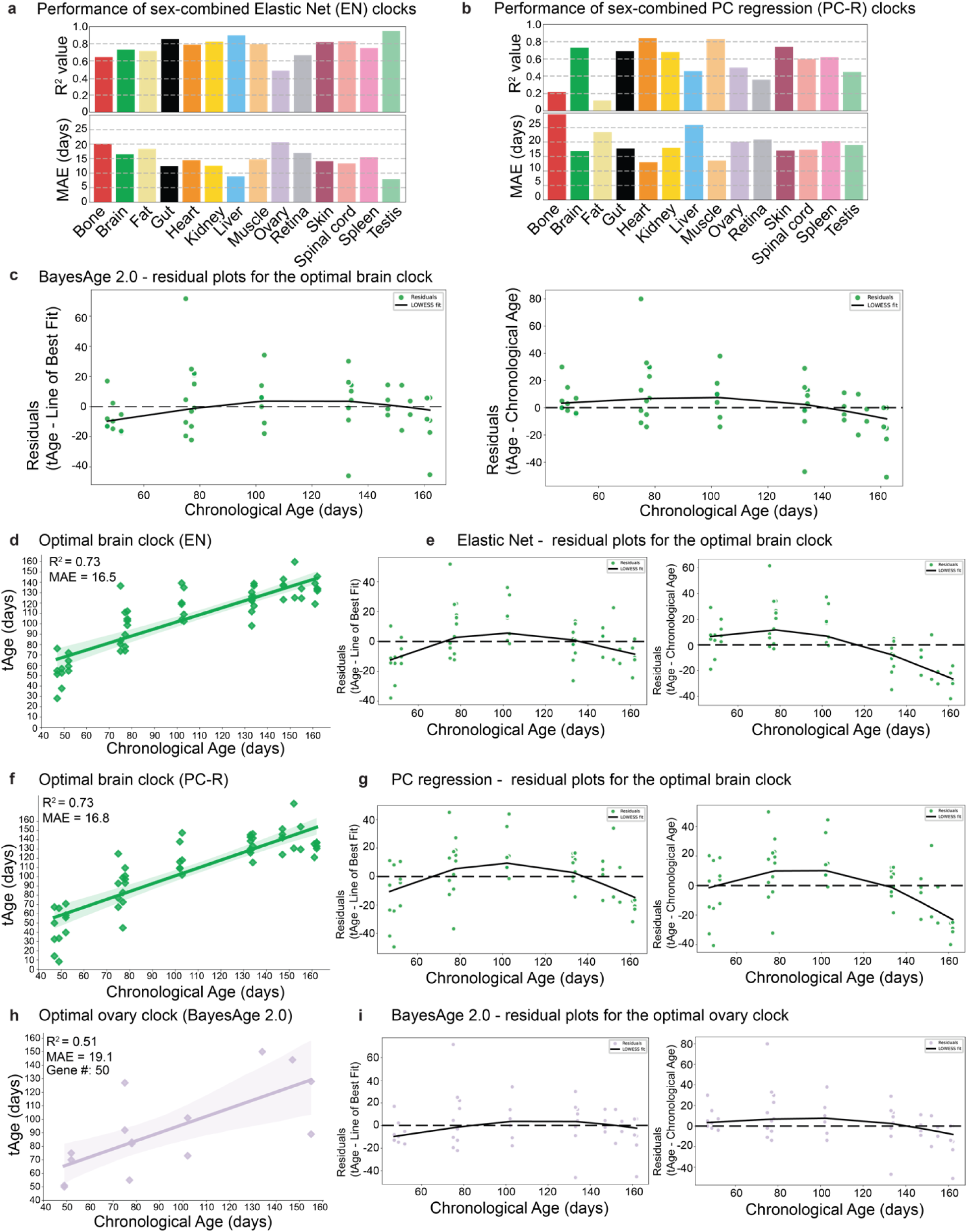
BayesAge 2.0 leads to less overfitting than Elastic Net and Principal Component regression models. (a-b) Bar plots of the performance metrics for (a) Elastic Net regression tissue clocks and (b) Principal Component regression tissue clocks, using the coefficient of determination (R^2^) for the relationship between chronological and predicted age and the mean absolute error (MAE). (c) Residual plots for the optimal brain clock modeled with BayesAge 2.0. Left, using difference between predicted transcriptomic age (tAge) and the line of best fit. Right, difference between predicted transcriptomic age (tAge) and chronological age. The ‘optimal’ BayesAge clock for a tissue is defined as the clock with the most concordance between chronological and predicted age. (d) Scatterplot of the tissue transcriptomic age (tAge) vs. chronological age for measuring the prediction accuracy of the optimal brain sex-combined tissue clock using Elastic Net regression. The coefficient of determination (R^2^) for the relationship between chronological and predicted age and the mean absolute error (MAE) are listed in graphs. The ‘optimal’ Elastic Net tissue clock is defined as the clock with the optimal combination of *α* and *λ* such that model error is minimized. (e) Residual plots for the optimal brain Elastic Net regression clock, calculated and plotted as in panel c. (f) Scatterplot of age predictions versus chronological age as in panel d for the optimal brain Principal Component regression (PC-R) clock. The ‘optimal’ PC-R tissue clock is defined as the clock with the optimal number of principal components such that there is the most concordance between chronological and predicted age. (g) Residual plots for the optimal brain PC-R clock calculated and plotted as in panels c and e. (h) Scatterplot of age predictions versus chronological age for the optimal ovary clock, the lowest performing tissue clock using BayesAge 2.0. (i) Residual plots for the optimal ovary BayesAge 2.0 clock, calculated and plotted as in panels c, e, and g.

**Extended Data Figure 7:**
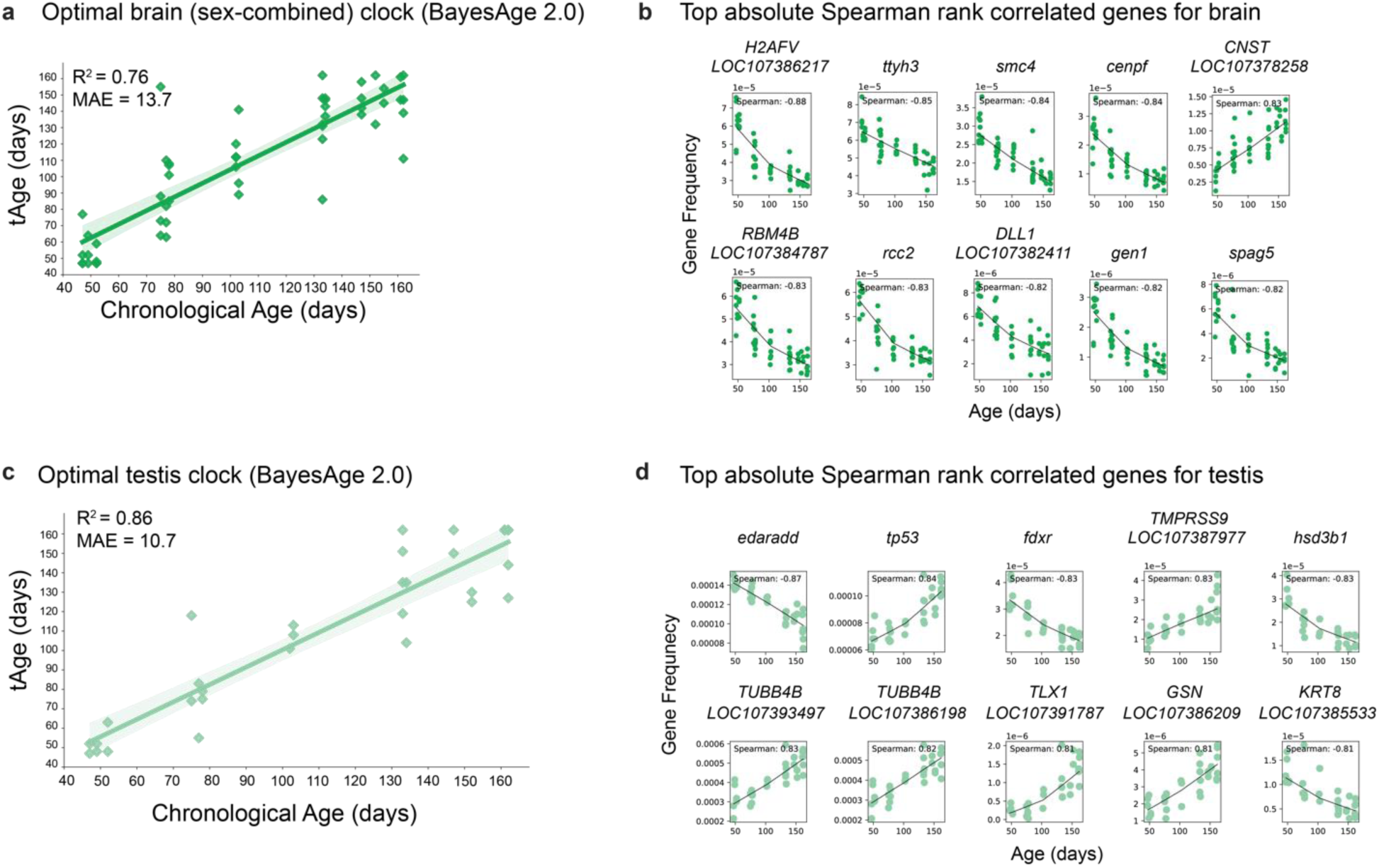
The brain and testis transcriptomic aging clocks are among the highest performing BayesAge 2.0 clocks across killifish tissues. (a) Scatterplot of the tissue transcriptomic age (tAge) vs. chronological age for measuring the prediction accuracy of the optimal brain sex-combined tissue clock, which is the model that corresponds to the most concordance between chronological and predicted age among all the gene number tested. The coefficient of determination (R^2^) between chronological and predicted age, as well as the mean absolute error (MAE), is listed in graphs. (b) The gene frequency scatterplots of the top 10 overall age-correlated genes trained on the sex-combined brain samples are shown. The black line is the locally weighted scatterplot smoothing (LOWESS) regression fit across time. (c, d) The scatterplots of tAge vs. chronological age (c) and gene frequency (d) were generated as in panels a and b, but for the testis.

**Extended Data Figure 8:**
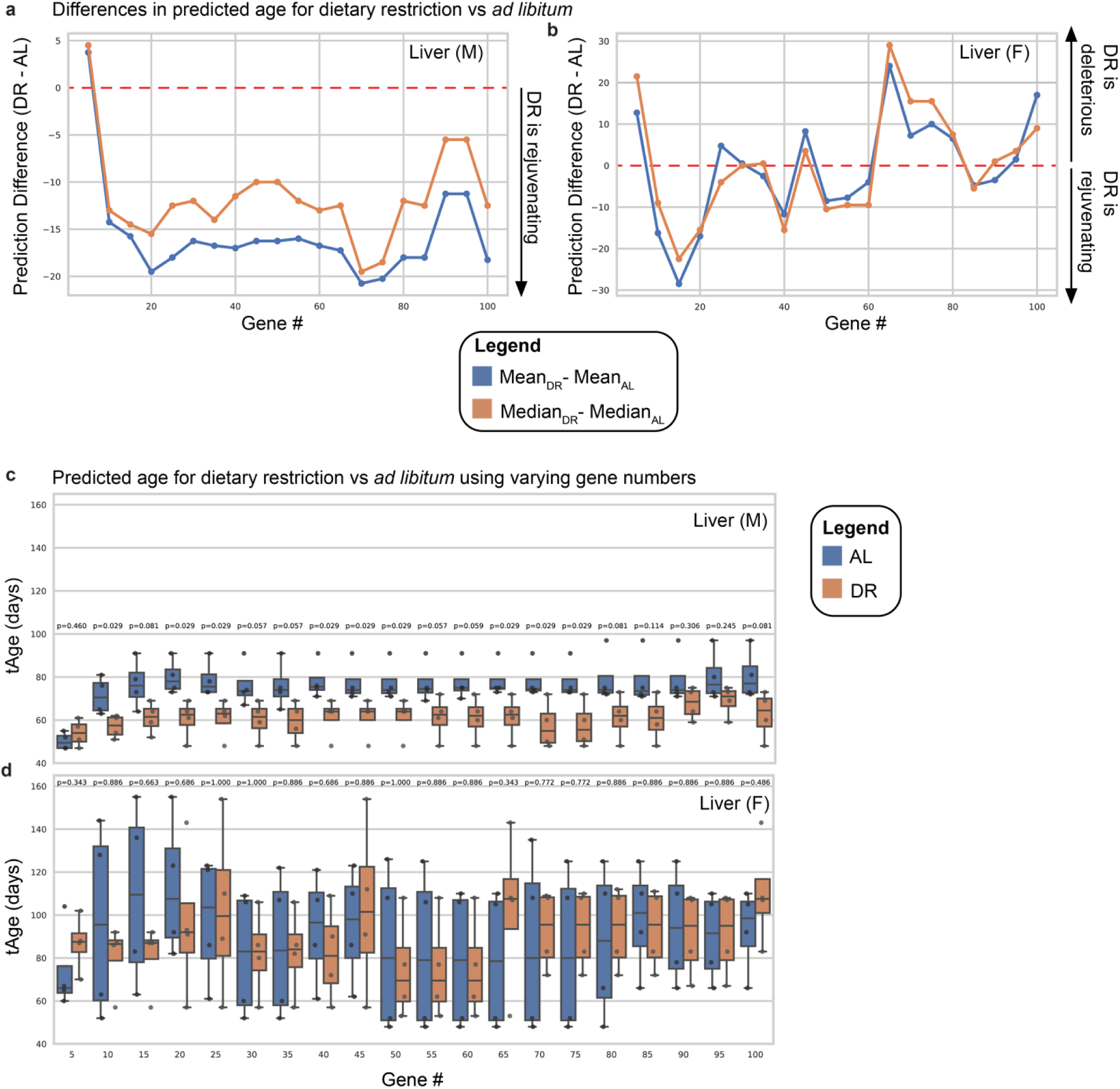
The sex-specific liver transcriptomic aging clocks predict dietary restriction results in ‘younger’ ages. (a, b) The predicted tAge difference between the *ad libitum* (AL) or dietary-restricted (DR) conditions observed across a range of clock gene numbers used for the male (panel a) or female (panel b) liver clocks. F, female; M, male. The median (orange) or mean (blue) predicted tAge was calculated from the 4 animals for each condition (AL or DR), and then the prediction difference in tAge was calculated by subtracting the median or mean in DR from that of the AL condition. Dotted line, AL and DR have the same predicted tAge. Below the dotted line indicates the DR condition is predicted to be ‘younger’ than the AL condition. The transcriptomic data were derived from a published dataset^22^. (c) Predicted tAges for the AL and DR conditions, male only, with each dot representing the predicted tAge of individual fish (4 fish per condition) when a specific clock gene number was used in the model. The box plots include the median, 25 (Q1), 75 (Q3) percentiles, and the whiskers include Q3+1.5×(Q3-Q1) and Q1−1.5×(Q3-Q1). At each gene number used for the model, Mann-Whitney test was used to test the significance of difference between the AL and DR conditions. (d) Predicted tAges for the AL and DR conditions, female only, plotted as described in panel c.

